# Residual flexibility in the topologically constrained multivalent complex between the GKAP scaffold and LC8 hub proteins

**DOI:** 10.1101/2024.11.25.624264

**Authors:** Eszter Nagy-Kanta, Zsófia E. Kálmán, Helena Tossavainen, Tünde Juhász, Fanni Farkas, József Hegedüs, Melinda Keresztes, Tamás Beke-Somfai, Zoltán Gáspári, Perttu Permi, Bálint Péterfia

**Affiliations:** Faculty of Information Technology and Bionics, Pázmány Péter Catholic University, Budapest, Hungary; Department of Biological and Environmental Science, University of Jyvaskyla, Jyvaskyla, Finland; HUN-REN Research Centre for Natural Sciences, Budapest, Hungary; Department of Chemistry, University of Jyväskylä, Jyväskylä, Finland; Institute of Biotechnology, Helsinki Institute of Life Science, University of Helsinki, Helsinki, Finland

**Keywords:** Postsynaptic density, protein NMR, molecular dynamics, PPI (protein-protein interaction), IDP (intrinsically disordered protein), multivalency

## Abstract

Guanylate kinase-associated protein (GKAP) is a large postsynaptic scaffold protein bearing two closely spaced noncanonical binding sites for the bivalent dynein light chain LC8 hub protein. This might allow the formation of heterogeneous complexes with different sizes and topologies. Here, we show that a well-defined hexameric complex is formed, composed of 2 GKAP molecules and 2 LC8 dimers. Using NMR spectroscopy, we demonstrate that the LC8-binding segment of GKAP is intrinsically disordered and the flexibility of the linker region is largely retained even in the complex form. Molecular dynamics calculations suggest that besides the tightly bound residues, the hexamer also exhibits several dynamically interchanging interactions. The flanking regions of the two binding sites on GKAP exhibit different interaction patterns, hinting at additional contacts that might explain the fixed stoichiometry of the assembly. Our results demonstrate that constrained stoichiometry can coexist with substantial flexibility in a multivalent system.

## Introduction

An increasing number of examples suggests that many intrinsically disordered proteins or regions (IDP/IDRs) are capable of multivalent binding and that the simultaneous formation of multiple interactions may be a common feature of IDPs [1]. There are many forms of multivalent assemblies, but the commonality is that a well folded domain participates besides an IDP or a disordered region [2]. Intrinsic disorder is a fundamental feature in the multivalent interactions formed by hub proteins. The ability to bind to a wide range of partner molecules makes hub proteins key components in maintaining the cell homeostasis [3]. The dynein light chain protein LC8 has been identified as a dominant, well-folded binding partner that can form multivalent complexes with many IDP/IDRs [2]. The two mammalian paralogs DLC1 (or DYNLL1) and DLC2 (or DYNLL2), both aliased as LC8, differ only in six out of 89 residues, and they are fully conserved as orthologs [4], [5]. LC8 was originally recognized as a member of the multi-subunit dynein motor complex, but it has also been described to interact with many partners in diverse cellular localizations. LC8 is considered a hub protein involved in many different cellular processes interacting with an extremely high variety of partners (more than 100 verified partners; *LC8 hub*, [6], Supplementary Table ST1). Many LC8 partners are primarily disordered [2] which makes LC8 a prime example of a multivalent folded protein where multispecificity is possible because of its preference for disordered partners (Teilum et al., 2021).

LC8 complexes are found in many different cellular assemblies [2], but the core structural basis for the complex formation is common [8]. Strong LC8 homodimerization (K_d_ ∼ 60 nM [4], [9], [10], [11] is necessary for the functionality, because both sub-units participate in the formation of the binding pocket. Disordered partners bind to the LC8 dimer through Short Linear Motifs (SLiMs). Motif preferences of LC8 binding partners have been described in the literature [12], [13], [14], [15]. Two loose consensus sequence classes have been proposed for LC8 partners: [K/R]_-3_X_-2_T_-1_Q_0_T_1_ and G_-2_I_-1_Q_0_V_1_D_2_ with the most conserved glutamine (Q_0_) set as the zero reference point. All known LC8 partners adopt a β-strand structure upon binding to the LC8 homodimer interface [16]. The LC8 interface contains a shared β-sheet with five strands (β1, β4, β5, β2 from one subunit, β3 from the other subunit), and a sixth strand is added by β-sheet augmentation from the binding partner [17], [18], while two α-helices stabilize this arrangement on the outer surface (α1 and α2) (Supplementary Figure S1). Additionally, the LC8 dimer is symmetric, therefore two identical binding sites are capable of interacting with two ligands, on the opposite sides of the dimer. Details of such interactions, involving both DLC1 and DLC2, are extensively described in the literature, almost exclusively based on LC8 complexes with small peptides. The majority of these contain one LC8 dimer, only a few higher-order systems have been studied so far (Supplementary Table ST1). All structures of such poly-multivalent complexes, where the LC8 binding partner contains multiple binding sites, have been solved by X-ray crystallography, where crystal contacts between different assemblies are unavoidably present, forming potential stabilizing interactions and likely constraining the observed complex geometries. The only exception is a study of the Chica protein by S. Clark and coworkers (BMRB ID: 26684, [19]), however, they only carried out NMR experiments with the free form of Chica with no NMR studies on the complex (Supplementary Table ST1). Thus, the flexibility and dynamics of multivalent LC8 complexes remains largely unexplored.

One of the many known LC8-partner molecules is GKAP (Guanylate kinase-associated protein, SAPAP1, Dlgap1). It is a postsynaptic protein interacting with several partners, including some of the most important proteins of the postsynaptic density (PSD) like PSD-95 and the Shank family [20], [21] (Figure 1A). GKAP is a scaffold protein contributing to the regulation, modulation and enhancement/amplification of signal transduction through the postsynaptic cell *via* NMDA receptors. GKAP is predicted to be largely disordered through its entire length, with high structural flexibility even in the binding regions, making it a perfect candidate to be an adjustable linker molecule in PSD [22]. GKAP harbors two LC8 binding sites (Figure 1B) and is likely to be able to bind multivalently to many other partners. The presence of multiple binding sites on GKAP have been experimentally confirmed for the proteins PSD-95 [23], Arc N-lobe [24] and Grb2 [25]. The available experimental structures for GKAP are limited to short peptides and its GH1 domain. This domain has been successfully crystallized and its structure consists of four α helices, however, its function is unknown [26].

**Figure 1.**
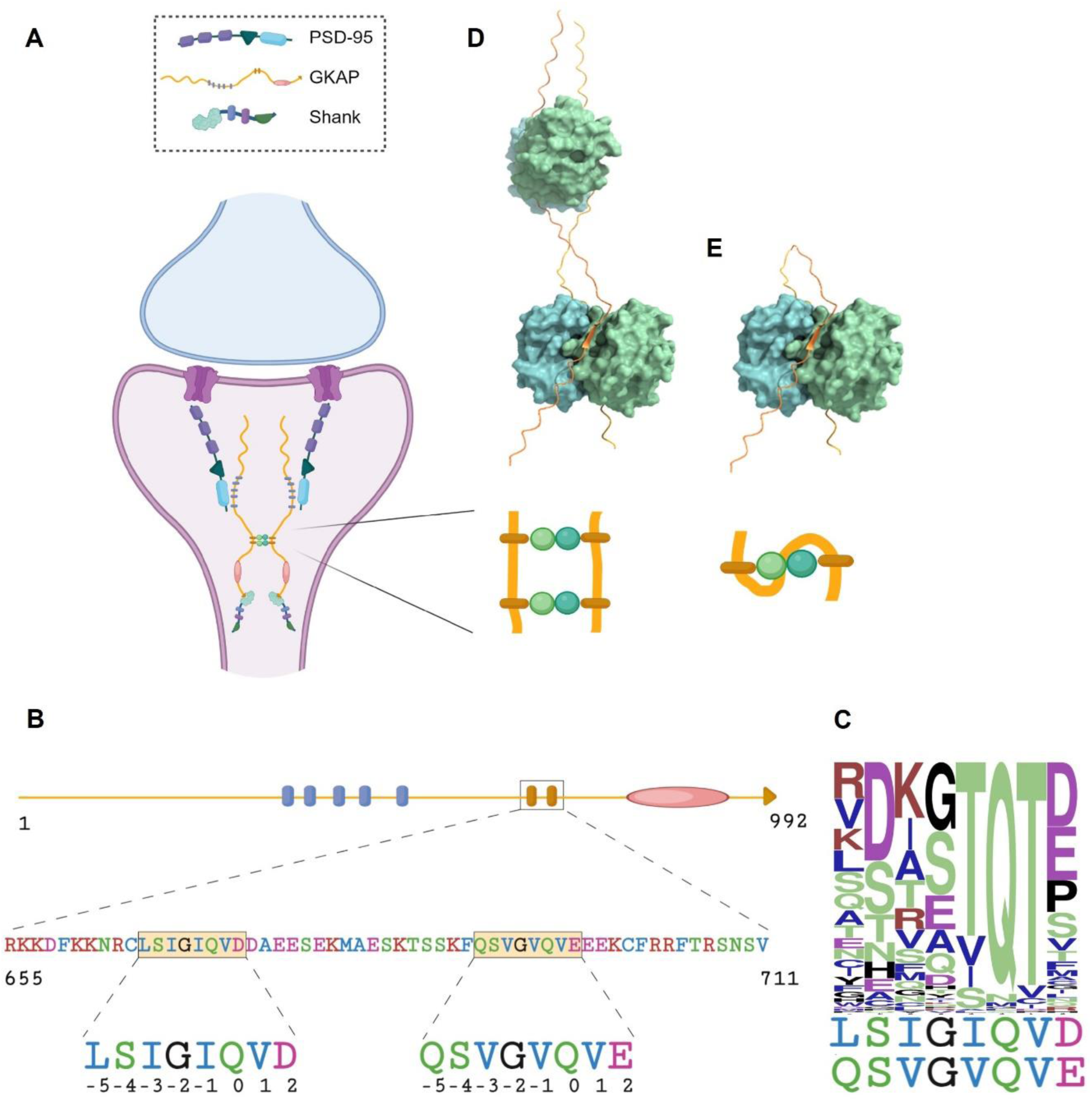
Interaction between GKAP and LC8 (A) GKAP is an abundant postsynaptic scaffold protein involved in the localization and organization of NMDA receptor complex. A plausible role of the interaction between GKAP and LC8 is to promote dimerization; (B) GKAP is predicted to be almost fully disordered with the exception of its C-terminal GH1 domain. Among others, GKAP contains five binding motifs for PSD-95 and two LC8 binding sites. The enlarged segment shows the construct used in this study with the two core LC8 binding motifs highlighted. The numbers refer to the positions within the binding motif, with relative numbering where Q0 is the anchor residue [13]; (C) Sequence logo of 117 LC8 binding motifs based on LC8 hub [34] and literature review (Supplementary Table ST1), with non-canonical binding motifs of GKAP displayed below the logo. Hydrophobic residues colored in blue, polar residues in green, acidic residues in magenta, basic residues in red, all other residues in black; (D-E) Proposed model of LC8:GKAP interactions from E. Moutin and coworkers with 2:2 or 1:1 stoichiometry (i.e. 2 GKAP: 2 LC8 dimers or 1 GKAP: 1 LC8 dimer)

The suspected role of the GKAP-LC8 interaction is in the organization of the NMDA receptor complex [27]. This interaction is suggested to be involved in the motor protein mediated trafficking, targeting and organization of the PSD-95 complex [27], [28]. GKAP as a scaffold molecule is a member of the core complex (PSD-95/GKAP/Shank complex) organizing glutamate receptors and the assembly of PSD [23], [27]. GKAP physically links NMDA receptors and surrounding scaffold molecules to the motor proteins of the cytoskeleton [26], [27].

The existence of the GKAP:LC8 interaction has been described and confirmed multiple times [12], [28], [29], [30]. The binding sites were specified with various methods, first with a yeast-two hybrid system [28], then later with pepscan technique [12], [30], [31] and co-immunoprecipitation [28], [29], [32]. The interaction has been experimentally investigated *in vivo* within a cellular environment using fluorescence fluctuation microscopy (two-photon scanning number and brightness, sN&B) [32] and live-cell confocal imaging [33]. Whereas DLC2 and DLC1 are highly related proteins having similar biochemical activities, based on yeast two-hybrid assays, GKAP seems to prefer DLC2 over DLC1 [28].

GKAP contains two linear motifs, having atypical amino acid composition, which are involved in LC8 binding. Both binding segments lack the consensus ‘TQT’ anchor region [34], and have atypical residues also in the other positions of the binding region (Figure 1B,C). The sequence logo shown in Figure 1C illustrates the frequent occurrence of the TQT motif in typical binding segments along with the diversity of the other positions in the motif. Nevertheless, the conserved binding mode of LC8 to its ligands (*via* ꞵ-sheet augmentation) suggests that its interaction with GKAP will also conform to this pattern. GKAP is capable of binding two LC8 dimers, and LC8 dimers are also able to bind to 2 ligands, therefore there is a possibility of forming polybivalent, hetero-oligomeric molecular scaffolds. In line with this consideration, E. Moutin and coworkers proposed two distinct types of organization for the GKAP-LC8 complex. They confirmed the complex formation *in vivo* [32], but the exact stoichiometry and the atomic structure of the complex is yet to be verified. Their hypothesis is that the complex is most likely formed with a 2:2 stoichiometry (*i.e.* 2 GKAP monomers and 2 LC8 dimers, Figure 1D,E).

In this study we describe the multivalent interaction between the LC8 dimer and the dynein binding region of GKAP as characterized by a combination of methods, including NMR spectroscopy. Our analysis provides a comprehensive description on the multivalent GKAP-LC8 interaction including NMR titration measurements in two, complementary setups: ^15^N, ^13^C labeled GKAP titrated with unlabeled LC8 and ^15^N, ^13^C labeled LC8 titrated with unlabeled GKAP. Our results suggest that predominantly hexameric complexes are formed (i.e. 2 GKAP chains binding 2 LC8 dimers), which, however, retain some of the flexibility characteristic of free GKAP, allowing the formation of several dynamically interchanging interactions.

## Results

### The LC8-binding segment of GKAP is disordered

For this study we selected the segment spanning residues 655-711 in *R. norvegicus* GKAP (UniProt ID: P97836-1) (Figure 1B). This region is 94.5% identical to the corresponding one in the human GKAP sequence (UniProt ID: O14490), with one substitution in the flanking, and seven substitutions in the linker region between the binding motifs. This segment, which we refer to as rGKAP_655-711_, contains both LC8 binding motifs with flanking regions of 10 residues on the N-terminus and 14 residues on the C-terminus.

The rGKAP_655-711_ construct exhibits typical properties of an extended IDP, namely, it shows large hydrodynamic dimensions, and it is characterized by low content of ordered secondary structure. The molecular weight (MW) calculated from the encoded amino acid sequence is 7 kDa (7014.88 Da), while on SDS−polyacrylamide gel electrophoresis (SDS−PAGE), the protein migrates abnormally slowly with an apparent MW of approximately 13 kDa (Figure 2A). The observed behavior of rGKAP_655-711_ in analytical size exclusion chromatography (SEC) also confirmed that its hydrodynamic dimensions are larger than expected for a folded protein. Based on the observed retention volume, the ratio of the measured vs. the theoretical molecular mass was 1.56, which corresponds to a completely unfolded state (Figure 2B, Supplementary Table ST2) [35]. The far-UV ECD spectrum of the construct was consistent with this (Figure 2C).

**Figure 2.**
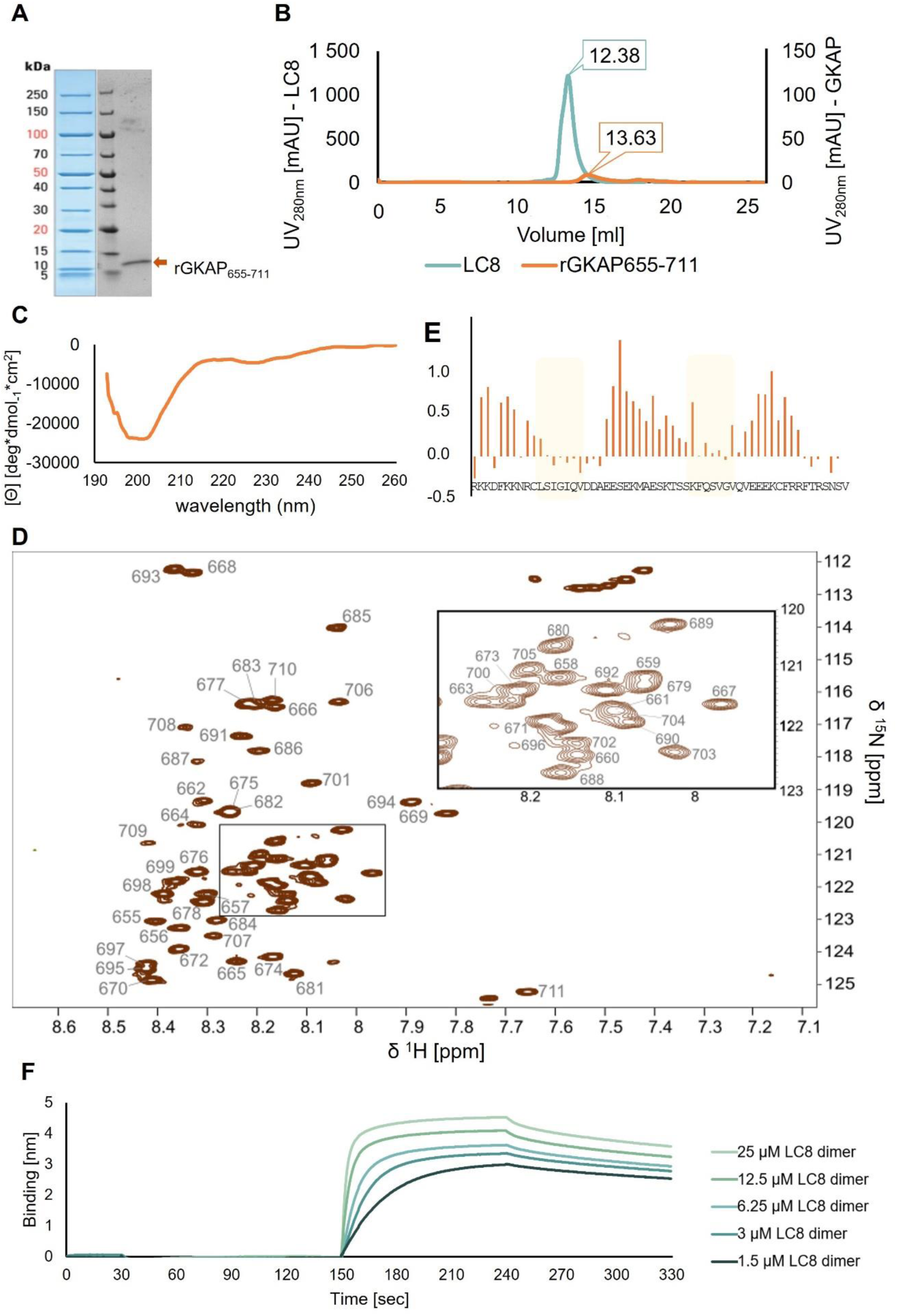
The LC8-binding segment of GKAP is disordered based on experimental data (A) On SDS-PAGE, GKAP migrates with an apparent molecular mass of 13 kDa instead of 7 kDa, as characteristic of disordered proteins; (B) Analytical size exclusion chromatogram of the rGKAP_655-711_ and the LC8 dimer for reference, showing larger hydrodynamic dimensions than in the case of an ordered protein (see Supplementary Table ST2); (C) Far-UV CD spectrum of rGKAP_655-711_ indicaties the absence of a well-folded structure; (D) ^1^H-^15^ N HSQC spectrum of rGKAP_655-711_ exhibits low signal dispersion, typical for disordered regions. Chemical shift assignments are denoted with the corresponding residue number. Inset shows expansion of the highlighted region between 120-123 ppm and 7.9-8.3 ppm for the ^15^N and ^1^H dimensions, respectively; (E) Sequential neighborhood-, temperature-and pH-corrected secondary chemical shift values shown as a difference for C^α^-C^β^ atoms indicate a disordered structure with slight preferences for extended structure in the binding regions and helical structures outside these; (F) BLI sensogram for the interaction of rGKAP_655-711_ and LC8, measurement of immobilized GKAP with free LC8.

Complete backbone and partial sidechain NMR chemical shift assignment of rGKAP_655-711_ (100% of the N, H^N^, C’, C^α^, C^β^, H^α^ and H^β^ atoms, and 52% and 61% of the C^γ^ and H^γ^ atoms, respectively) was deposited in BMRB. The C^β^ chemical shifts of both cysteines (C664 and C701) were 27.9 ppm, consistent with the presence of free thiol groups in the buffer containing TCEP.

Signal dispersion in the ^1^H, ^15^N HSQC spectrum of the rGKAP_655-711_ construct is ∼1.2 ppm in the ^1^H and ∼13 ppm in the ^15^N dimensions, respectively (Figure 2D). Sequential neighborhood-, temperature-and pH-corrected secondary chemical shift values [36] for C^α^-C^β^ atoms are between-0.2 and +1.2 ppm (Figure 2E). Secondary chemical shift values for the H^α^ atoms is below 0.13 ppm (Supplementary Figure S2). These observations confirm that the construct is intrinsically disordered along its full length. The calculated differences between C^α^-C^β^ secondary chemical shifts indicate a slight preference for extended conformation in the first LC8 binding motif (Figure 2E, residues 667-674) and varying helical propensity outside the LC8 binding motifs (residues 656-665, 675-688 and 696-704). CheSPI [37] analysis also shows a mainly disordered structure with no strong preference for stable secondary structure formation, based on the C’, C^α^, C^β^, N and H^N^ chemical shifts (Supplementary Figure S3).

### GKAP binds LC8 at both binding sites simultaneously

Biolayer interferometry (BLI) experiments show that LC8 binding to GKAP follows pseudo-first-order kinetics with a single transition (Figure 2F). The dissociation constant K_d_ of the complex is 0.29 μM (R^2^=0.9779, for aspecific binding see Supplementary Figure S4). This implies a relatively strong interaction compared to the nNOS peptide binding to LC8 dimer (5.41±0.15 μM measured with Isothermal titration calorimetry, ITC), another protein with a non-canonical LC8 binding motif (having IQV instead of TQT residues in the central region [38], Supplementary Table ST1). Association rate constant (k_a_) is 8800 ± 83 M^-1^s^-1^, while dissociation rate constant (k_d_) is 2.5 ·10^-3^ ± 4.3 · 10^-5^ s^-1^.

NMR titration of ^15^N, ^13^C-labeled rGKAP_655-711_ with unlabeled LC8 (referred to as “forward” titration henceforth to distinguish this setup from the “reverse” titration where the labeling was the opposite, see below, Figure 3A-B) revealed the involvement of both LC8 binding sites on GKAP. We monitored binding using ^1^H-^15^N-HSQC at GKAP:LC8 ratios of 1:½, 1:1, 1:2, 1:4 (considering the dimeric concentration of LC8). There are residues at both binding sites with clearly disappearing amide N-H peaks starting at an estimated molar ratio of 1:1 (equal to 2 GKAP with 2 LC8 dimer) during titration, with no sign of heterogeneity at the population level. In other words, LC8 seems to bind equally tightly to the two GKAP sites. The last two titration points (GKAP:LC8 dimer 1:2 and 1:4) are virtually indistinguishable indicating that saturation was reached (Figure 3A). The limited number of peaks and the position of peaks implies sample homogeneity in the titration endpoints. The observed behavior of the peaks can be either due to the formation of a high molecular weight species or extensive line broadening due to exchange phenomena occurring at μs-ms timescale. In either case the simplest interpretation is that the corresponding residues participate in interactions that are not present in the absence of LC8. We note that very similar behavior is observed in the reverse titration for the amide peaks of ^15^N, ^13^C-labeled LC8 when complexed with unlabeled rGKAP_655-711_, strongly suggesting the formation of a molecular species largely invisible for NMR (Figure 3B). Results from dynamic light scattering (DLS) experiments are consistent with a largely homogeneous fraction of the GKAP:LC8 complexes formed. The estimated hydrodynamic diameter is in the range of ∼5 nm, larger than the estimates obtained for the unbound rGKAP_655-711_ segment and the unliganded LC8 dimers (∼1-4 nm and ∼3-4 nm, respectively). This approximated diameter is also consistent with the calculated molecular weight of ∼61 kDa of a hexamer containing four LC8 chains (two dimers) and two rGKAP_655-711_ segments (Supplementary Table ST3). Furthermore, Far-UV CD spectrum of the GKAP:LC8 complex is slightly different from the sum of the spectra of the individual partners, consistent with the presence of the interaction but not indicative of any large-scale structural rearrangement in the partners (Supplementary Figure S5).

**Figure 3.**
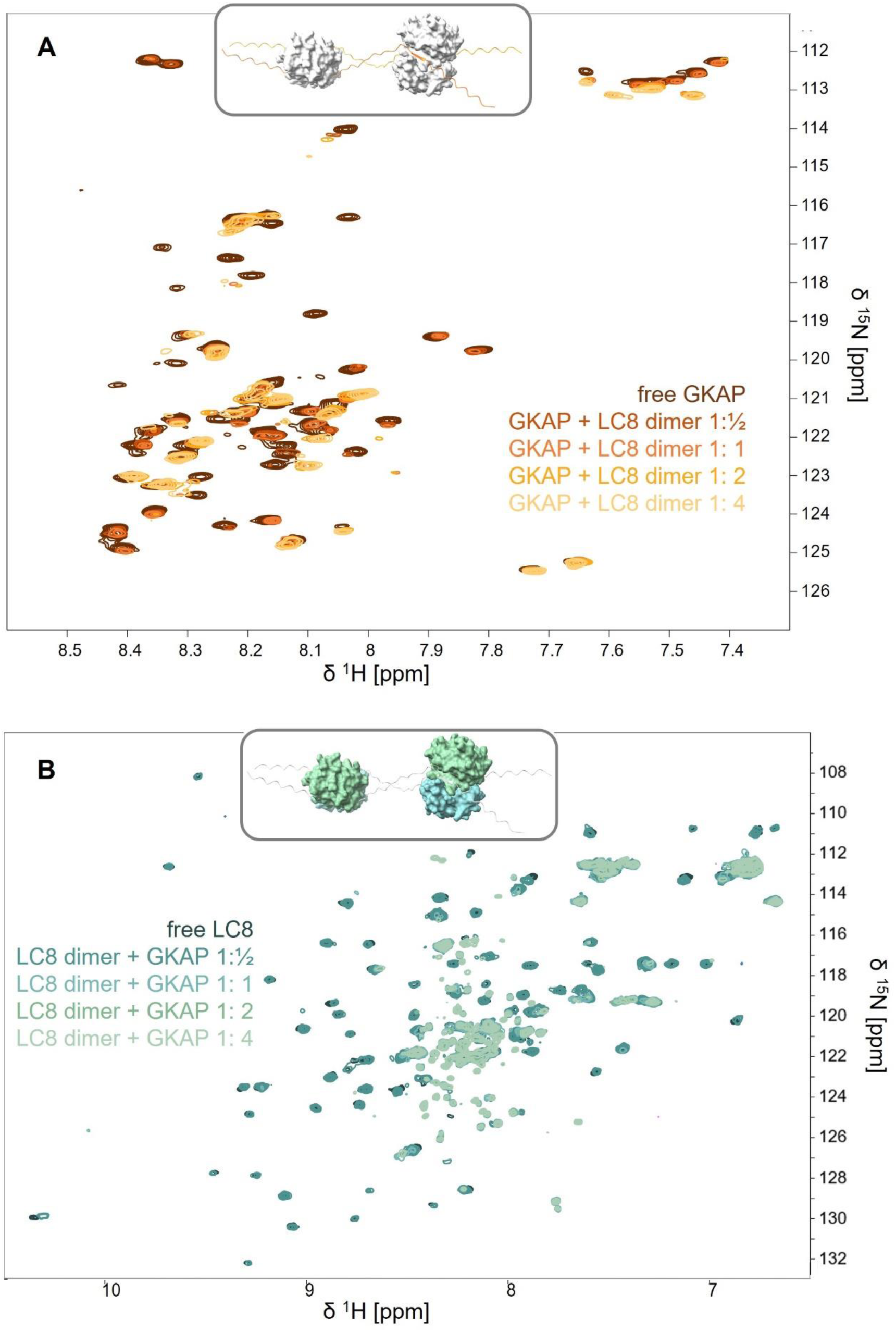
NMR titration experiments monitoring the GKAP:LC8 interaction. (A) “Forward” titration: ^1^H-^15^N-HSQC of GKAP in the presence of LC8 dimers at ratios of 1:½, 1:1, 1:2, 1:4. (B) “Reverse” titration: ^1^H-^15^N-HSQC of LC8 dimers at GKAP ratios of 1:½, 1:1, 1:2, 1:4; Insets show a structural model of the hexameric complex with the ^15^N-labeled components colored according to the titration setup.

These observations suggest that the emerging complex has a well-defined, hexameric stoichiometry with two GKAP segments bound by two LC8 dimers. The complex also presumably has a preferred topology in terms of the arrangement of the binding sites, meaning that one LC8 dimer binds to identical sites on the two GKAP segments, as proposed in Figure 3C by E. Moutin and coworkers [32]. Structural modeling reveals that the 17-residue linker between the two LC8 binding sites enables the simultaneous binding of two LC8 dimers along each GKAP segment arranged in a parallel geometry. Dimensions of our hexameric structural models fall in the range of the experimentally determined size of the complex (Supplementary Table ST3). Our results rule out the possibility of the 1:1 stoichiometry (or 2:1 according to E. Moutin and colleagues [32], Figure 1DE), where the two binding motifs of one GKAP molecule interact with one LC8 dimer, and the GKAP disordered linker between the binding motifs is wrapped around the compact dynein dimer.

### The central region of GKAP remains disordered in the complexed state

Curiously, a number of peaks remain virtually unchanged in the ^1^H-^15^N HSQC spectra of ^15^N-labeled rGKAP_655-711_ complexed with unlabeled LC8. The majority of these peaks can be mapped to residues in the region between the two LC8 binding sites, indicating that this part of the polypeptide largely retains its disordered nature even in the complex form (see N and H^N^ chemical shift perturbation data on Figure 4A). Consequently, the linker region between the two binding sites on rGKAP_655-711_ also retains much of its flexibility, and its chemical environment is not considerably different in the bound and the free form.

**Figure 4.**
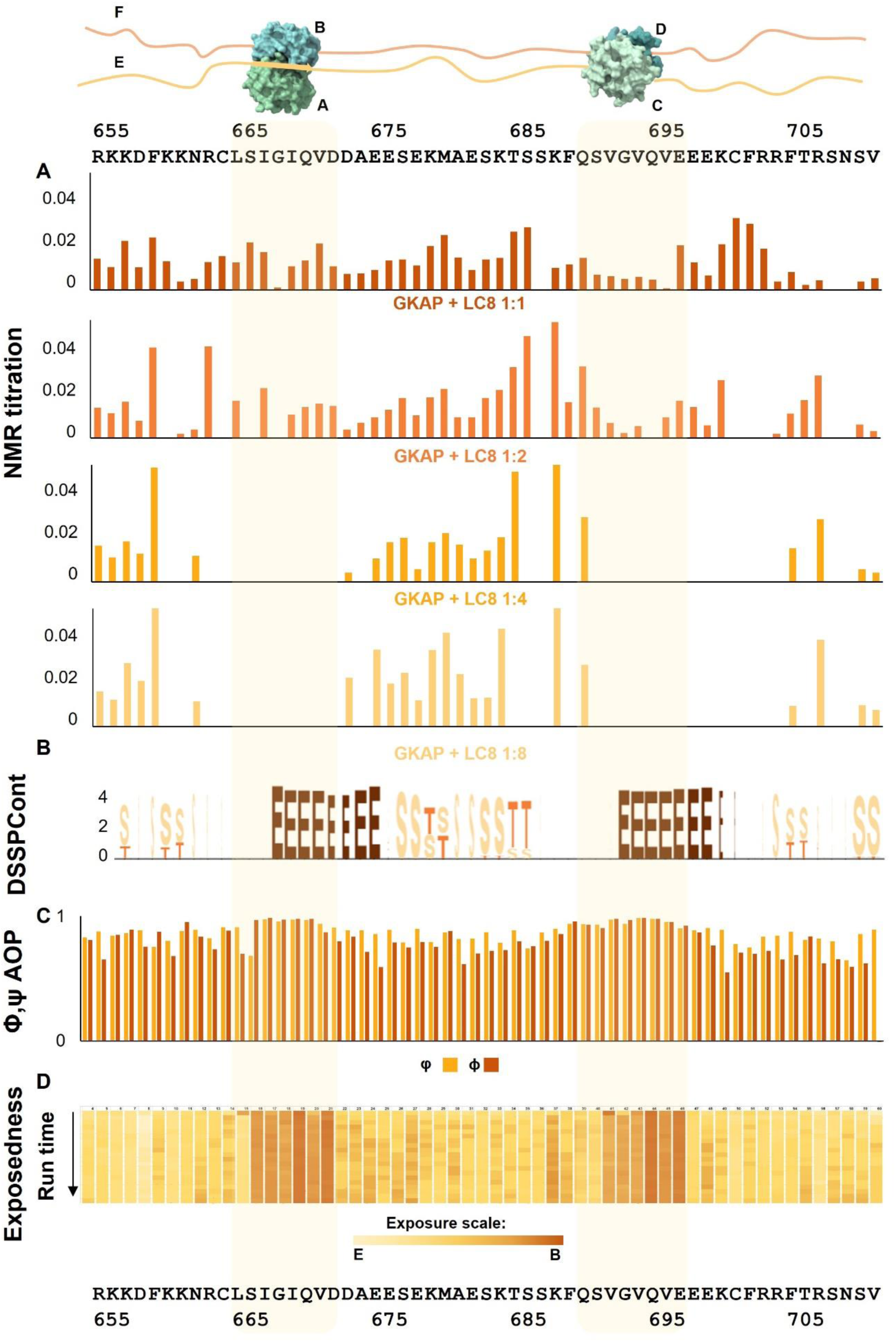
Structural properties of LC8-bound GKAP (A) Chemical shift perturbation data of the “forward” titration points, calculation: 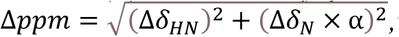where α=0.14; (B) Representative DSSPCont example showing secondary structure elements from one GKAP chain in a selected molecular dynamics run of the complex (for all chains see Supplementary Figure S7); (C) Angular order parameters (AOP) of the backbone φ and ψ angles calculated for the one GKAP chain in a selected MD run of the complex; (D) Changes in solvent exposure during the simulations of the complex as calculated by DSSP (average of the MD runs for the manually built models).

All amide peaks disappear from their original positions in the neighborhood of the first binding motif, from K_-10_ to A_4_, with the exception of N_-8_ and D_3_ (see the relative amino acid numbering in Figure 1B). All the peaks disappear in the neighborhood of the second binding motif, from T_-10_ to N_14_, with the exception of K_-7_, Q_-5_, F_7_, and R_9_. Although intensive signal overlap and strong ambiguity can be observed in the ^1^H-^13^C HSQC titration spectra (Supplementary Figure S6), the persistence of disappearance of several peaks can be confirmed even when considering their C^α^-H^α^, and C^β^-H^β^ peaks. The following residues undoubtedly disappear from both ^1^H-^15^N HSQC and ^1^H-^13^C HSQC spectra, with both the C^α^-H^α^, and C^β^-H^β^ peaks: C_-6_, L_-5_, I_-3_, G_-2_, I_-1_, Q_0_, D_2_ in the first, and F_-6_, G_-2_, V_-1_, Q_0_, V_1_, E_4_, C_6_, R_9_, T_11_ in the second binding motif. Besides this, several peaks on the N terminus and in the linker region between the binding motifs remain visible in the same position with approximately the same intensity during the titration steps as in the free form (Figure 4A).

Based on molecular dynamics (MD) simulations, in the bound form the two binding sites in GKAP form a β-strand structure, but they behave slightly differently, especially when also considering their flanking regions. DSSP [39] cannot consistently assign regular secondary structural elements to a significant fraction of amino acid residues outside the binding region (in some cases it assigns helices). With DSSPcont (continuous secondary structure assignment, [40]) the main characteristic of the linker seems to be flexible/disordered, however a slight preference for helical conformation is observable in some of our simulations, which is in line with experimental results (Figure 4B).

Our MD simulations along with AlphaFold Multimer-based modeling suggest that the relative orientation of the two LC8 dimers in the hexameric assembly can be highly variable. It is consistent with the presence of a flexible linker between the binding sites (Figure 5A). However, in our simulations, the typical scenario shows the two LC8 dimers approaching each other and forming contacts, with minimal changes in their relative orientation throughout the remainder of the simulations. In this respect, both these and the various models obtained with AlphaFold Multimer represent largely static snapshots, rather than offering direct insight into the dynamic rearrangement that is plausibly the scenario most consistent with our experimental observations. The retained structural flexibility is also confirmed by the angular order parameters calculated for the backbone φ/ψ angles of GKAP conformers in the MD trajectories. The values are consistently lower outside of the binding regions, indicating higher structural variability (Figure 4C).

**Figure 5.**
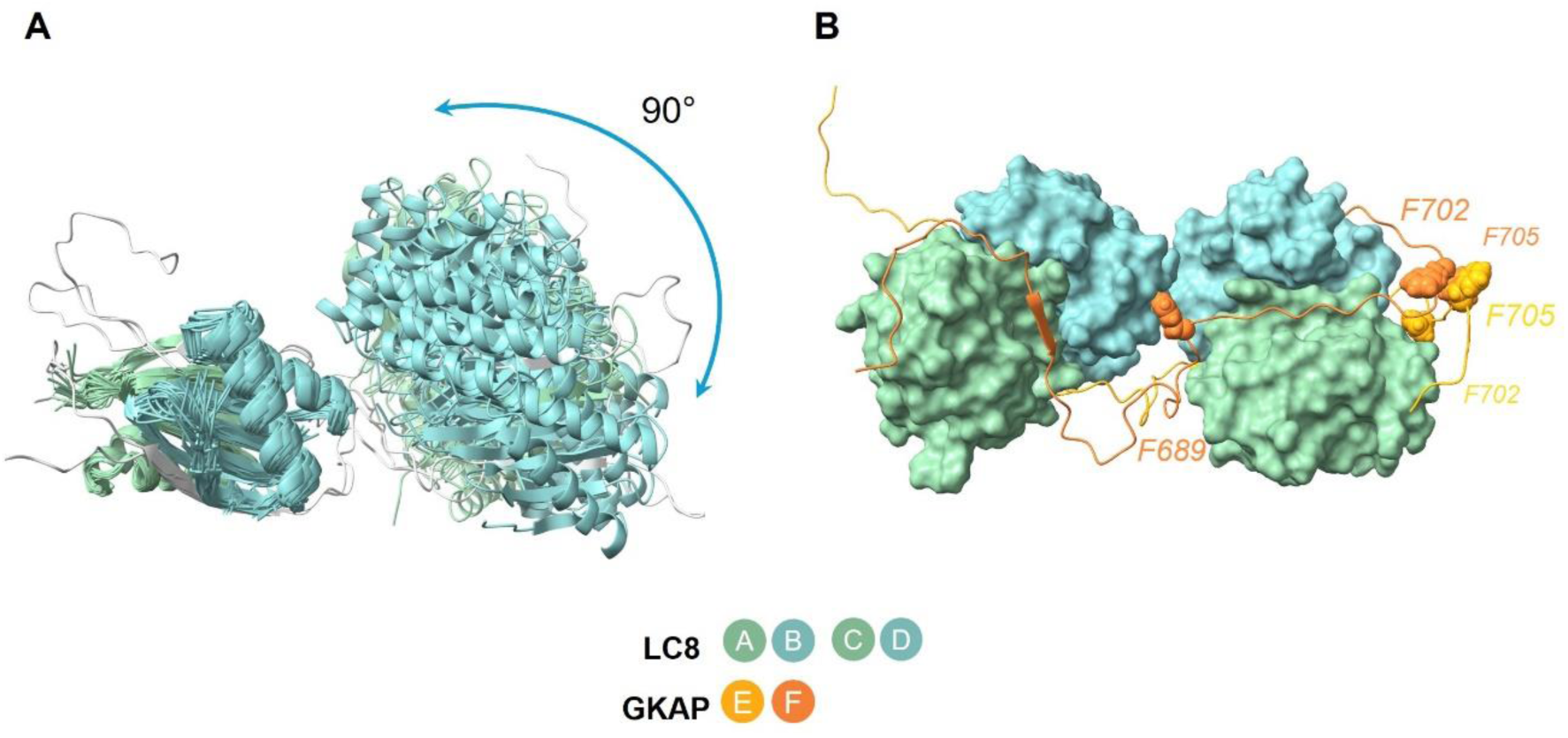
Molecular dynamics results highlight the disordered nature of GKAP (even in complex with LC8). (A) high flexibility, possibility of LC8 dimer-LC8 dimer contact; (B) Phenilalanine residues (F689, F702 and F705, highlighted with sphere representation) raising the possibility of GKAP-GKAP contact.

### The two GKAP binding sites show a slight difference in LC8 binding

NMR data show that both GKAP binding sites are affected during binding, however, there are differences between their structural properties. For both binding sites the affected peaks reveal the involvement of residues outside the immediate binding site, up to position-10 towards the N-terminus and at least to position 4 towards the C-terminus. The flanking regions around the binding sites participate in the structural stabilization of the interaction, but in different ways (Supplementary Figure S8). The peaks corresponding to the flanking regions surrounding the binding motifs mainly disappear (with the exception of 2-4 residues) from the ^1^H-^15^N HSQC titration spectra (Figure 4A), but with slightly different patterns for the two sites. Two explanations are possible for this phenomenon I) either these regions are wrapped around the LC8 dimer, losing their flexibility, or II) the incorporated amino acids participate in intermediate exchange time-scale events.

In the MD simulations, two lysine residues in positions-10 and-9 of the N-terminal binding region (K660 and K661) seem to be able to form interactions with LC8, but these are not stable, and, most importantly, it is apparently not possible for both residues to be bound simultaneously. The most likely explanation for this is that no backbone conformation is compatible with both side chains being buried. Our hypothesis is that these interactions are dynamically interchanging at the μs-ms time scale, causing line broadening and thus rendering the peaks “invisible” for NMR. No characteristic difference is observed between the core IQV and VQV binding motifs either in the NMR spectra or in the MD simulations: all corresponding peaks disappear at 1:4 concentration (along with other neighboring residues) and they also exhibit similar residue-residue contacts in the trajectories. The common feature is that positions-1 and +1 are extremely buried and make contact with LC8 residues forming strands β3 and β4 (Figure 4D).

Positions-8,-7 and-5 exhibit different behavior in the two binding sites, despite belonging, in some cases, to chemically similar amino acids. For example, in position-7, the amide peak of R663 in the first binding site completely disappears, whereas the peak of the corresponding K688 in the second site shows a moderate position shift. The simulations show that these positions are buried over time in the first binding site and maintain significant interactions with LC8. The N-terminal flanking region of the first binding motif participates loosely in the interaction. F689 in position-6 with regard to the second binding site can either contact the LC8 dimer at the second site or that binding to the first, N-terminal site. The observed flexibility of the linker region suggests that this residue might not be tightly locked in a single position, rather undergoing binding and unbinding events (Figure 5B). Even in the binding motif, a possible fast exchange phenomenon can be observed in the-4 position serine, but due to lack of bound-form assignments this cannot be unambiguously concluded.

The differences are even more pronounced for the C-terminal flanking segments, with residues in positions 5-14 affected only in the second, C-terminal binding site, most of them disappearing completely (Figure 4A). The corresponding positions relative to the first N-terminal binding site coincide with the linker between the sites and are only moderately affected, indicative of segments remaining largely disordered as described above. These differences reveal the importance of investigating constructs with longer flanking regions in contrast with studies focusing only on the minimal binding regions. Our MD calculations suggest that this region could potentially be buried and come into significant proximity to LC8, but no consistent pattern is observed, in line with the experimentally observed dynamic nature of the region. The most remarkable difference between the two binding sites is that the C-terminal binding site and its C-terminally located flanking region seem to be involved in complex formation *via* possible exosite contacts. While in the case of the N-terminal binding motif, residues immediately after the minimal binding site, starting from position +3-with the sole exception of A_4_-give rise to N-H peaks in the HSQC spectrum of the complex, indicating residual flexibility, residues after the C-terminal site become undetectable by NMR. This kind of difference cannot be readily explained by our MD simulations, where both flanking segments appear partially buried (Figure 4D). However, it has been noted previously that LC8 bears a positively charged surface patch where it interacts with these positions [19], [41], and thus the different arrangement of negatively charged residues, especially the series of three glutamates at positions 2-4 after the C-terminal site, might contribute to the observed differences.

In several simulated conformers, the two GKAP segments make extensive hydrophobic contacts with each other using two phenylalanine side chains, F702 and F705, in the C-terminal flanking region after the second binding site. This suggests that the two GKAP chains might form specific interactions with each other, facilitated by the binding of the LC8 proteins (Figure 5B, Supplementary Figure 9). It might be speculated that this interaction might serve as a trigger for the formation and stabilization of an otherwise weak and noncanonical coiled coil segment.

### The two LC8 dimers interact with each other in the hexameric assembly

We have performed a partial backbone assignment of the LC8 dimer. Although multiple data sets for LC8 chemical shifts are available in BMRB, none of the available assignments were performed in a TCEP-containing buffer. Reverse titration, using ^15^N, ^13^C-labeled LC8 and unlabeled rGKAP_655-711_ revealed profound changes in the ^1^H-^15^N-HSQC spectrum of LC8. At the monomer LC8:GKAP ratios of 1:½ (equaling GKAP:LC8 dimer ratios of 2:2, the stoichiometry we confirmed, where hypothetically all LC8 dimers are in bound form) and at titration points with higher excess of GKAP, most LC8 peaks disappear. However, many peaks in the middle ^1^H region of the spectrum are still present. We investigated the system further by recording sensitive, overnight ^1^H-^15^N spectra, in which peaks belonging to both the ^15^N-labeled LC8 along with those of the unlabeled GKAP were observed. These latter peaks are located in positions matching the remaining peaks in the forward titration, i.e. the ^1^H-^15^N-HSQC spectrum of ^15^N-labeled GKAP in complex with unlabeled LC8. Moreover, ^15^N-filtered ^1^H spectra clearly indicate that these peaks arise from ^1^H-^14^N groups, i.e. unlabeled GKAP, in the reverse titration experiments. This confirms the disappearance of the majority of LC8 ^1^H-^15^N peaks (Figure 3B, Supplementary Figure S10).

The remaining LC8 peaks include those corresponding to the amide NH of residues D3, R4, A6 and S88, which can be unambiguously assigned. A few more peaks also clearly belong to the LC8 dimer, however, based on the acquired spectra they could not be reliably assigned (unfeasibility of 3D experiments of the complex). Our MD calculations indicate that there are some residues, mainly in helices α1 and α2, that are not involved in either dimerization or ligand binding (Supplementary Figure S11). On the other hand, MD simulations suggest that a multitude of interactions might be formed between the LC8 dimers binding to different binding sites. These can involve different residues at the N-terminus, helices α1 and α2, the strand β1 and also loop regions. Thus, it is likely that the behavior of some the amide peaks belonging to residues outside the ligand binding site can be explained by intermolecular interactions between the two LC8 dimers, while unaffected peaks again indicate residual flexibility also in this partner. Residues R4 (near the N terminus) and N51 in different LC8 dimers are brought into physical contact in one of our MD trajectories. This might be due to the fact that the N-terminal region of the LC8 chains remains slightly flexible, also confirmed by the NMR “reverse” titration experiments.

The general observation is that peptide binding does not significantly change the conformation of LC8 apo-form [8]. Our MD simulation of LC8 indicates that the structure of LC8 dimers is not significantly affected by ligand binding.

## Discussion

Our NMR and CD data clearly demonstrate the intrinsically disordered nature of the LC8-binding segment of GKAP. Thus, the LC8:GKAP interaction involves two kinds of bivalent partners, the globular LC8 dimers and the highly flexible GKAP segments. The behavior of the NMR peaks in both the forward and reverse titrations along with DLS and SEC-based size estimates points to the emergence of a single species with a large molecular weight that is largely “invisible” to NMR investigation. In contrast, specific regions both on GKAP and LC8 retain considerable flexibility and still give rise to observable peaks in NMR, and the different behavior of peaks together with MD simulations hint at the presence of potential dynamic rearrangements within the complex. MD simulations are expected to yield insights into the behavior of the system. In our case, because of the scarcity of NMR data on the complex, no quantitative agreement can be enforced between experiment and simulation. It cannot be necessarily expected that a conventional simulation recapitulates all structural and dynamical aspects of the complex, thus, we have chosen to explore the MD trajectories whether any of them can provide clues for selected observations. Some general phenomena are consistently reproduced by the simulations (e.g. retained flexibility between binding sites, see Figure 5A) however, many unique events and interactions can be observed only occasionally, at different points of the separate runs.

The significance of the disappearing peaks can be evaluated in the light of the previously published literature on the structures of LC8 and its ligands. Interestingly, the first (N-terminal) binding region of GKAP has a L in position-5. According to Rapali and collegues, in natural binding partners this position seems not to favor any specific residue type [14]. In contrast, in their *in vitro* evolutional study there is a preference for apolar (mainly V, but also M/I/L) or long aliphatic (R/K) amino acids and this results in increased affinity. In the second (C-terminal) binding region, position-5 is occupied by Q which does not belong to any of the aforementioned categories (Supplementary Figure S8). Serine in position-4 is a shared feature between the two binding motifs, and besides backbone hydrogen bond forming, it might also participate in sidechain hydrogen bonding based on our results and literature data [14], [19], [42].

Based on previous studies, the residues from positions-8 to +4 in the LC8 binding motif are relevant for the binding affinity in a monovalent interaction with LC8 [12], [43]. In the case of short peptides representing the minimal binding motif, the presence or absence of a residue in the-5 position can change the dissociation constant by ten-to hundred-fold [14]. Our results emphasize the relevance of the regions flanking the binding motif (from-10 to +4, and even to +14 in the case of the second binding motif), refining the results from previous efforts to characterize the minimal sequence segments required for binding (LSIGIQVD, QSVGVQVE, from-5 to +2). Our most intriguing finding is that while the linker between the two sites retains its flexibility, a remarkably long C-terminal flanking region of the second binding motif is involved in the interaction.

Like GKAP, the nNOS peptide is an IQV motif partner. By performing a mutational analysis on nNOS (-6 MKDTGIQVDRDL +5) it was found that mutating the two hydrophobic amino acids flanking the central glutamine residue to asparagines (i.e. Ile in-1 and Val in +1 position) completely abolishes the interaction. They found that I57, F62, F73, A82, L84 and F86 of LC8 interact with these two amino acids [8], which is consistent with our MD results. On the “reverse” NMR titration spectra I57, L84 and F86 disappear from the spectra (F62 and F73 cannot be assigned).

The C-terminal flanking region of the second binding motif participates in the complex formation. Negatively charged residues at the +2 and +3 positions are able to form electrostatic interactions with spatially close charged amino acids of LC8 [19], [41]. The PDB structure 1F96 is the only available NMR structure for a ligand bound LC8 dimer. In this case, a longer C-terminal region was part of the construct. Based on the deposited NMR distance restraints, Gln in +3 and Gln in +5 are in contact with residues in the β1-β2 loop and the strands β3 and β4 of LC8 (Supplementary Figure S1). Our observations are in line with these results, thus, it is feasible that residues following the core binding motif also form interactions with LC8 but this might largely depend on the actual amino acid sequence.

One of the main features of the LC8 hub protein is that it forms polymultivalent interactions by binding to multiple motifs, which in most cases involves two consecutive binding sites on their partner molecule. In general, it seems that the canonical “TQT” motif containing partners bind LC8 with the highest affinity. As the sequence diverges from this pattern, binding affinity decreases (resulting in higher K_d_). As multivalent interactions are usually formed to strengthen the generally weak binding mediated by SLiMs, it is plausible to speculate that the presence of consecutive non-TQT motifs result in higher overall affinity compared to that of a single motif, possibly due to avidity. K_d_ values (Supplementary Table ST1) of the LC8 interactors clearly indicate an increase in binding strength in favor of partners with two binding motifs over single binding motifs, by at least one order of magnitude. Our results (K_d_=0.29 μM) are in line with this trend. For the GKAP-LC8 hexameric complex, the observed K_d_ is smaller than for the similar “IQV” nNOS peptide (sequence: DTGIQVD, K_d_ = 5.4 μM) and the association and dissociation rate constants also show a higher binding affinity (for nNOS peptide, k_a_ = 46800 M^-1^s^-1^ and k_d_ = 4.37 · 10^-1^ s^-1^) [38].

In LC8 partners, the two LC8-binding sites are usually located ∼20 amino acids apart. The GKAP:LC8 interaction also follows this pattern. The LC8 hub database [34] lists 14 bivalent or multivalent LC8-interacting proteins. LC8 was proposed to act as a dimerization engine, and some of its interaction partners, like Swallow was validated [44] and others were predicted to undergo oligomerization *via* coiled-coils, with examples including the proteins Ana2, Bassoon, Kibra, Nup159 [15], [45], [46], [47]. For Ana2, the coiled-coil region is located between the two binding sites [15].

Our systematic search for multivalent protein:LC8 interactions resulted in five similar structures in the PDB, with a longer protein segment (27-33 residues) incorporating two distinct LC8 binding motifs in complex with two LC8 molecules. Two of them (PDB ID: 3FM7 [48] and 2PG1 [49]) are complexes of one DLC1 dimer, one DLCTctex-type dimer and two disordered dynein intermediate chains having two LC8 binding sites. The PDB structure 3GLW is a quasi-hexamer generated *via* translational and rotational transformations of a DLC1 monomer and an intermediate chain partner peptide unit [48]. The fourth structure contains DLC1 dimers with the protein Panoramix (PDB ID:7K3J [50]) and the fifth is a DLC1-SAO-1 complex (PDB ID:7Y8 [51]) (Supplementary Table ST1). All these PDB structures were solved by X-ray crystallography and all these complexes have an elongated shape with a parallel topology, meaning each LC8 dimer either binds two N-terminal or two C-terminal sites on the partners. This arrangement is consistent with our model based on NMR and other data. In the X-ray structures, the linker regions between the LC8 binding sites adopt an extended conformation, and the LC8 dimers binding to the same partners typically do not interact with each other (crystal contacts between different hexamers are possible). However, these features might be the results of crystal packing, and the flexibility of these segments cannot necessarily be directly assessed based on these structures.

Here we provide experimental evidence that the linker region between the two binding motifs remains highly flexible even when both binding sites are occupied and the hetero-hexamer is formed (Figure 5A). In the structures mentioned above, 3-9 residues are located between the two binding motifs, while in GKAP the linker region consists of 17 amino acid residues. In addition, none of the LC8 partner proteins in the solved structures have amino acid compositions similar to that of GKAP in their binding motifs (harboring the canonical TQT or TQV anchor sequence in the binding motif, except for SAO-1, having VAT and CQT).

Analysis of the linker regions between LC8 binding motifs was performed by the group of E.J. Barbar using the multivalent IDP ASCIZ having seven LC8 binding motifs [3], [52]. They conclude that shorter linker regions cause increased rigidity of the resulting complexes. Based on their proposal, the longer linker region resulting in higher flexibility might explain the functionality of the complex as an assembly-scaffold, a regulator-modulator of the postsynaptic density signal transduction pathway initiated at NMDA receptors [52]. They also proposed a classification of linker lengths as a short, a long, and a “mid-length” in bivalent/multivalent LC8 partners [3]. According to this, GKAP is a mid-length bivalent ligand, and here we propose that this results in the formation of a single well-defined complex with LC8.

The Barbar group also investigated the Chica-LC8 interaction, where four binding motifs are present on the Chica protein. The Chica protein was proved to be intrinsically disordered by NMR, but solution-state studies of its complexes with LC8 were hampered by solubility issues. Structures of LC8 with peptides corresponding to individual binding motifs have been solved by X-ray crystallography (PDB IDs: 5E0L, 5E0M [19]). The LC8 binding motifs on the Chica protein show clearly distinct sequence patterns, also leading to differences in their binding properties. They found that the K_d_ value of the multivalent binding (when 4 binding motifs are included in the sequence) equals the K_d_ value belonging to the peptide binding with the strongest affinity, and the multimer with multiple bound LC8 dimers does not show enhanced stability. It was proposed that the presence of additional binding sites promotes self-association. We note that this scenario also leads to a topologically constrained complex, where the positionally equivalent LC8 binding sites are bound by the same LC8 dimer on both partner molecules, an arrangement consistent with our observations for the GKAP:LC8 interaction. Although there is no coiled coil predicted with high confidence in GKAP, the output of COILS [53] with a 21-residue window size hints the presence of a weak or non-canonical short coiled coil between residues 715-735 (towards the C-terminal end of the second binding motif) (Supplementary Figure S12). It might be speculated that LC8 binding might trigger the formation of a more extended interaction between GKAP monomers, possibly even forming a short coiled coil, consistent with the behavior of several other LC8 partners and the dimerization promoting role of LC8. This kind of interaction between the GKAP molecules could also provide an explanation for the preferred stoichiometry and arrangement of the hexameric assembly, as well as the observed binding kinetics.

Our results demonstrate the large added value when combining the power of NMR with molecular dynamic calculations to study complex, flexible systems. The unexpected revelations related to the role of the flanking regions in the interaction highlights the importance of investigating protein segments including all binding motifs and their neighboring regions instead of short peptides. Examining the residue-level connections between the postsynaptic density scaffold GKAP and the hub protein LC8 forms the basis of understanding their role and mechanism of action in the postsynaptic density. The flexibility of the hexameric system described here in detail might indicate their functionality as an assembly scaffold or modulator-regulator complex in the NMDA receptor-initiated signal transduction pathway in the postsynaptic cells.

## Methods

### Lead contact

Further information and requests for resources and reagents should be directed to and will be fulfilled by the lead contact, Bálint Péterfia.

### Experimental details

#### Protein expression and purification

The GKAP construct was designed to include both LC8-binding motifs of GKAP with extended flanking regions (10 residues on the N-terminus, 14 residues on the C-terminus). The segment spanning residues 655-711 in the *Rattus norvegicus* GKAP isoform 3 “GKAP1a” (UniProt ID: P97836-5) was selected. The insert was picked up from a cDNA (ORF template) kindly provided by Enora Moutin. The insert was amplified for cloning and ligated into NdeI and XhoI sites of an altered pEV vector (Novagen) that contains an N-terminal 6xHis tag and a tobacco etch virus (TEV) protease cleavage site. The actual construct contains four extra residues (GSHM) at the N-terminus, remaining from the expression tag.

The pEV plasmid vector construct containing the *Rattus norvegicus* DYNLL2 gene (UniProt ID: Q78P75, 100% identical to the human ortholog, UniProt ID: Q96FJ2) was kindly provided by Prof. László Nyitray. This pEV vector also contains an N-terminal 6xHis tag, the TEV protease cleavage site and four residues (GSHM) at the N-terminus remaining from the expression tag.

Both protein constructs were produced in BL21 (DE3) *E. coli* (Novagen) cells. Cells were transformed with the vectors described above and grown in LB media, or in minimal media containing ^13^C-labeled _D_-glucose and ^15^N-labeled NH_4_Cl as the only carbon and nitrogen source, respectively. Protein expression was induced with 1 mM IPTG (Isopropyl β-D-1-thiogalactopyranoside) at 6 MFU cell density and the recombinant proteins were expressed at 20°C overnight. Centrifuged cell pellets were stored at-20°C until further usage.

Cell pellets were lysed by ultrasonic homogenization in 10% cell suspension using a lysis buffer (50 mM NaPi, 300 mM NaCl, pH 7.4). In the case of rGKAP_655-711_, denaturing-renaturing IMAC purification was applied. After homogenization, cell pellets were dissolved in denaturing buffer (6 M GdnHCl, 50 mM NaPi), added to 5 ml Nuvia™ Ni-affinity column (Bio-Rad), then proteins bound to the column were renatured with native buffer (50 mM NaPi, 20 mM NaCl, pH 7.4). After a washing step with 50 mM, elution was performed with 250 mM imidazole, and was followed by His-tag removal with TEV protease. For LC8, the same purification protocol was used but after ultrasonic homogenization and centrifugation, the supernatant was immediately purified with IMAC Nuvia Ni-affinity column.

Protein samples were concentrated by ultrafiltration using Amicon® Ultra Centrifugal Filter with 3 kDa molecular weight cut off value, and the buffer was changed to low salt NaPi Buffer (50 mM NaPi, 20 mM NaCl, pH 6.0). Proteins were further purified by ion exchange chromatography, using 5 ml High Q column with the same buffer (50 mM NaPi, 20 mM NaCl, pH 6.0; recombinant proteins were collected in the flow through fraction). After another step of protein concentration, proteins were further purified with size exclusion chromatography (SEC) on a Superdex™ 75 Increase 10/300 GL 24 ml column, the buffer was changed to 50 mM NaPi, 20 mM NaCl, pH 6.0. Later 0.02% NaN_3_, and 5 mM TCEP (pH 7.4) was added to every sample individually.

The concentration of LC8 was measured by its absorbance at 280 nm using a NanoDrop2000 photometer, while the concentration of GKAP was measured with Qubit Protein assay. The recombinant proteins’ purity and exact molecular weight was analyzed by SDS-PAGE and LC-MS. The SDS-PAGE gels were evaluated with the software GelAnalyzer (GelAnalyzer 23.1.1 (available at www.gelanalyzer.com) by Istvan Lazar Jr., PhD and Istvan Lazar Sr., PhD, CSc).

#### Analytical size exclusion chromatography (SEC) measurements

In the analytical SEC measurements, the standard proteins: BSA, myoglobin, RNAse A, B12 vitamin were dissolved in the same buffer as GKAP and LC8 (50 mM NaPi, 20 mM NaCl, pH 6.0, 5 mM TCEP) in 200, 2500, 3000 and 300 µM concentration, respectively. Proteins were injected one by one in the same volume (500 µl) as GKAP and LC8 to a Superdex™ 75 Increase 10/300 GL 24 ml column, and the flowspeed was between 0.8-1 ml/min.

#### Far-UV CD spectroscopy

Proteins were measured by ECD spectroscopy using JASCO J-1500 spectrometer (JASCO Corporation, Tokyo, Japan). ECD spectra were recorded at 20 °C using 0.1 cm path length J/21 quartz cuvette (Hellma absorption cell) and 300 µl sample with the following settings: 195– 260 nm spectral range, 50 nm/min scanning speed, 1 nm bandwidth, 0.2 nm step size, 0.5 s response time and 3 scans of accumulation and baseline correction. rGKAP_655-711_ was diluted to 6.5 µM in 50 mM NaPi, 20 mM NaCl, pH 6.0 buffer containing 5 mM TCEP. Free LC8 was measured in the very same buffer, diluted to 11.25 µM concentration (considering monomers). The complex was created with the same molar concentrations (i.e. 6.5 µM concentration of GKAP with 11.25 µM concentration of LC8). Thus, we expect that the majority of the molecules is present as part of the complex, which is the dominat species under the conditions used. Comparison of the CD spectra was performed under the assumption that the signal solely represents the complex, and it should be noted that even if this is not completely true, this does not affect our conclusions obtained from the qualitative analysis as described.

#### Biolayer interferometry

Prior to experiments, all Ni-NTA biosensors (Fortebio, USA) were hydrated in kinetic buffer (50 mM NaPi, 20 mM NaCl, pH 6.0 + 20 mM imidazole, 0.1% BSA, 0.02% Tween 20, 0.02% NaN_3_) at 25°C for 10 min. The ligand, the 6xHis-tagged rGKAP_655-711_ protein was immobilized on the surface of the biosensor at a 5 µg/ml concentration in kinetic buffer for 90 sec. The initial baselines were then recorded in kinetic buffer for 30 s. The association and dissociation sensorgrams of LC8 dimers at concentrations ranging from 3.125 μM to 50 μM were recorded in kinetic buffer for 90 sec each. The equilibrium dissociation constant (K_d_) was determined from the BLI data using the global fitting method provided in the data analysis software. The curves were adjusted with a background spectrum (with 0 added ligand). To check for aspecific binding, the experiment was repeated with the same settings but without immobilizing the 6xHis-tagged rGKAP_655-711_ construct on the sensor surface in the first step. During evaluation of the aspecific binding experiment, a baseline was measured with kinetic buffer only.

#### Dynamic light scattering

Particle size and size distribution of GKAP, LC8 dimer and their complex were measured at 20 °C using a W130i dynamic light scattering device (DLS, Avid Nano Ltd., High Wycombe, UK) with a diode laser (660 nm) and a photodiode detector. 80 μl samples were used in low-volume disposable cuvettes with 1 cm path-length (UVette, Eppendorf Austria GmbH). The time-dependent autocorrelation function was measured for 10 s, repeated 10 times. Data analysis was performed using i-Size 3.0 software.

#### NMR measurement conditions

NMR experiments were acquired using 0.06–0.5 mM ^15^N, ^13^C-labeled protein samples in 2/98% D_2_O/H_2_O at pH 6.0. Chemical shifts were referenced to external 2,2,-dimethyl-2-silapentane-5-sulfonic acid (DSS). All data were acquired at 298.15 K on a Bruker AVANCE III HD 800 MHz spectrometer, equipped with a TCI ^1^H/^13^C/^15^N Z-gradient cryoprobe. Data was collected and processed with TopSpin version 3.5.7.

The following experiments were used in the resonance assignment of rGKAP_655-711_: ^1^H-^15^N HSQC, constant-time ^1^H-^13^C HSQC-aliphatic, constant-time ^1^H-^13^C HSQC-aromatic, HNCACB, HN(CO)CACB, HNCO, HBHA(CO)NH, (H)CC(CO)NH, H(CCCO)NH [54], [55] and i(HCA)CO(CA)NH [56]. In the case of LC8, partial backbone assignment of the ^1^H-^15^N-HSQC was performed mainly in comparison with bmrb15076 [41], with the support of HNCACB and HN(CO)CACB spectra. This was needed for the evaluations of the titration ^1^H-^15^N HSQC spectra, to identify the disappearing and visible peaks corresponding to the free LC8 peaks. NMR titration experiments were conducted with the unlabeled partner molecules at titration points visible on Figure 3.

CCPNmr Analysis v3.1 software was used for the chemical shift assignment.

#### Model building and MD simulations

For the initial model building PDB ID: 2XQQ [14] was used as a template. The extended disordered regions were modeled using Dipend and Modeller (Chimera) [57], [58]. Another set of models were generated using AlphaFold in the complex modeling setting in the Colabfold notebook [59]. FoldX was applied before MD simulation.

Computations were performed on the Komondor supercomputer. The models were simulated with GROMACS [60] using explicit solvent classical MD. AMBER99SB-ILDN protein force field was used; the box setting was a minimum distance of 1.0 n. For the initial model five parallel 200 ns runs were carried out (md_200_1, md_200_2, md_200_3, md_200_4, md_200_5). The AlphaFold generated models each had a 200 ns run time (AF1, AF2, AF3, AF4, AF5). From the ten 200 ns long production run 10 x 20 000 snapshots were taken.

The MD generated models were analyzed as follows: R Secondary structure was predicted using DSSP and DSSPcont consecutively [39], [61]. Exposure was calculated from DSSP. Radius of gyration was calculated with and in-house script. φ/ψ angles were calculated for each model using phipsi.py script from UCSF and further analyzed with an in-house script. For all other calculations and file formatting we used Python 3 codes written in our group. Coiled-coil prediction was made using wagga-wagga webserver (10.1093/bioinformatics/btu700). The most representative peptide was collected using MolMol [62].

The residue level molecular interactions were analyzed using Voronota [63], from which the average atomic distances were obtained. For visualization we used ChimeraX, Weblogo and Jalview [64], [65], [66].

## Acknowledgements

We are grateful for Elnora Moutin for providing the rGKAP cDNA, and for László Nyitray for providing the rDYNLL2 vector construct. We would like to thank László Nyitray for his valuable insights regarding LC8. We would like to thank Viktor Farkas for the opportunity to perform ECD measurements, the training and the guidance in the evaluation of the results. We would like to thank Laszlo Dobson for his help with Voronota. We are grateful for the Jyväskylä Team’s help in the sample preparation for the experiments: Ilona Pitkänen and Maarit Hellman.

We are thankful for all the current and former lab members for providing additional help, namely Zita Harmat, Brigitta Maruzs, Anna Sánta, Mária Stenbach and Soma Varga.

This work was supported by the Hungarian National Research, Development and Innovation Office (NKFIH) (grants OTKA NN 124363, OTKA K 137947, TKP2021-EGA-42 to B.P. and G.Z.). This work was also supported by the Jane and Aatos Erkko Foundation and grants from the Academy of Finland (grant 323435 and 362535 to P.P.).

## Author Contributions

Conceptualization, Z.G.; Methodology, Z.G., B.P., T.J. and P.P; Investigation, E.N-K., Zs.E.K., H.T., T.J., F.F., J.H., M.K. and P.P.; Writing – Original Draft, E.N-K., Zs.E.K. and Z.G.; Writing – Review & Editing, E.N-K., Zs.E.K., H.T., T.J., T.B-S., P.P., B.P., and Z.G.; Supervision, P.P., H.T., B.P., T.J. and Z.G.; Visualization, Zs.E.K. and E.N-K.; Funding Acquisition, Z.G.; Resources, B.P., P.P. and G.Z.

## Supplementary Figures

**Supplementary Figure S1:**
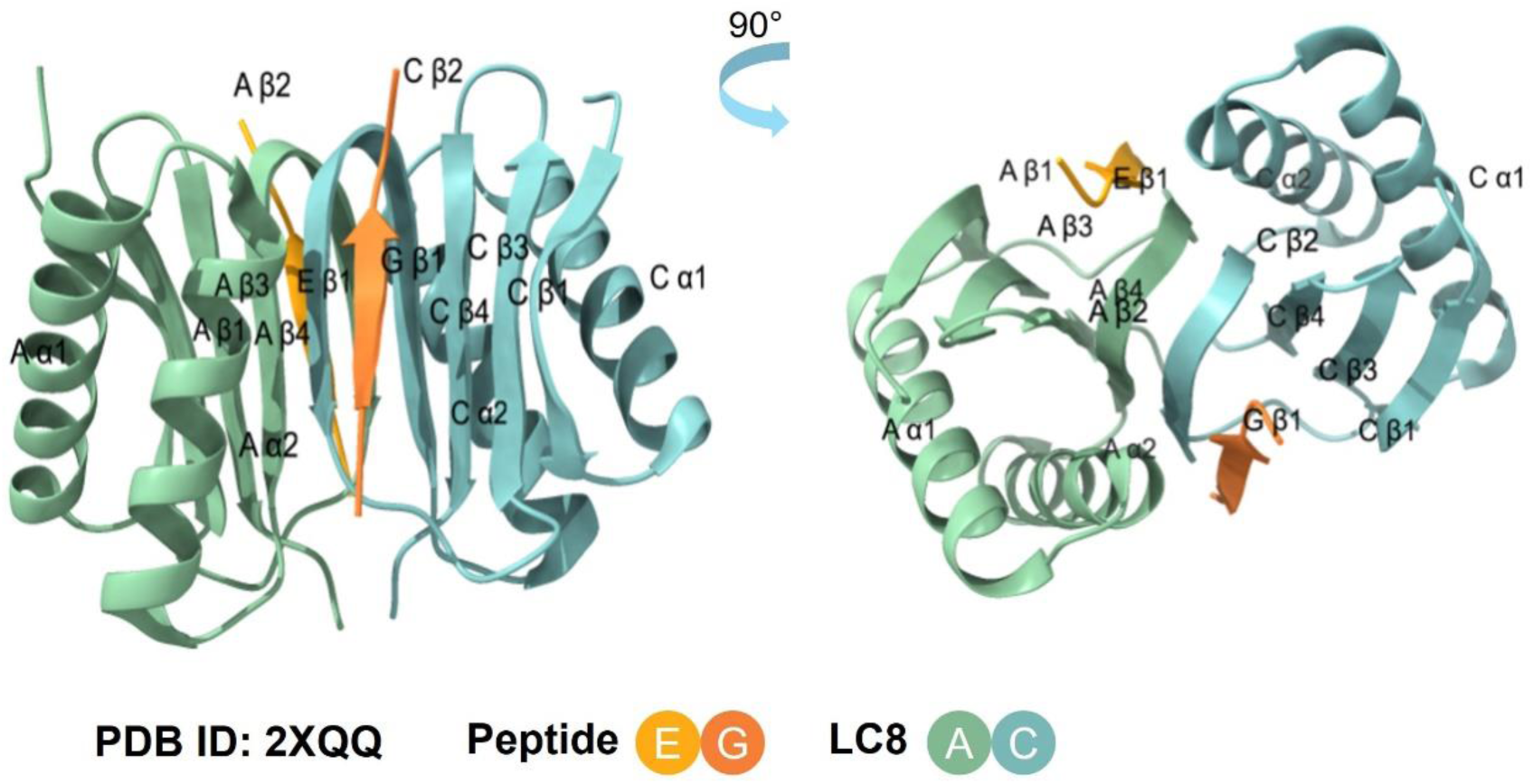
Common nomenclature of LC8 dimer secondary components. This nomenclature is used throughout the article. (Representation PDB ID: 2XQQ)

**Supplementary Figure S2.**
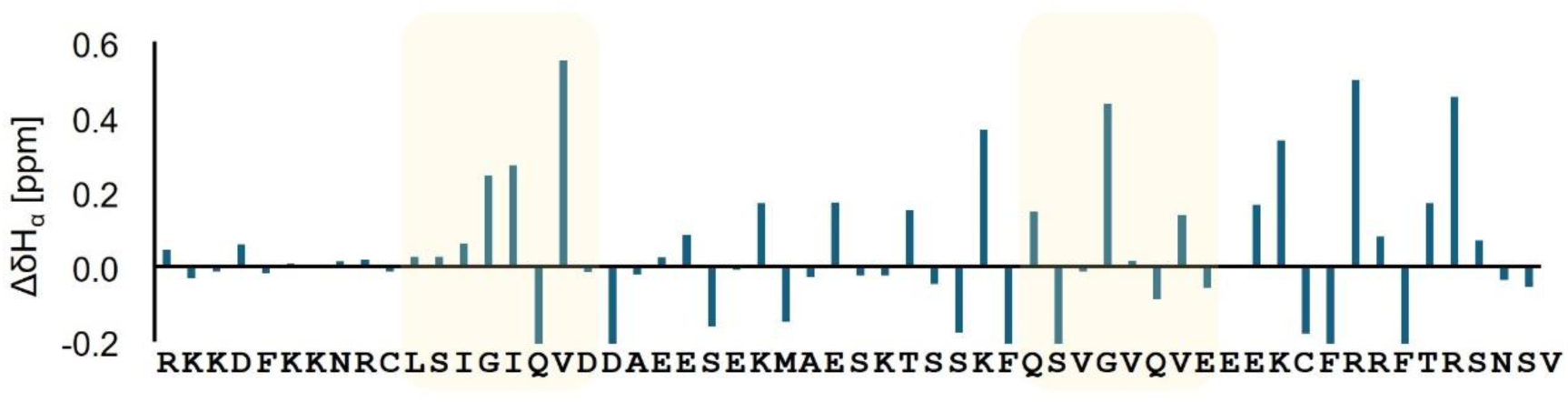
Sequential neighborhood-, temperature-and pH-corrected secondary chemical shift values for the H^α^ atoms [36]. Calculated with pH 6.0, 298 K and ionic strength of 0.07 M.

**Supplementary Figure S3.**
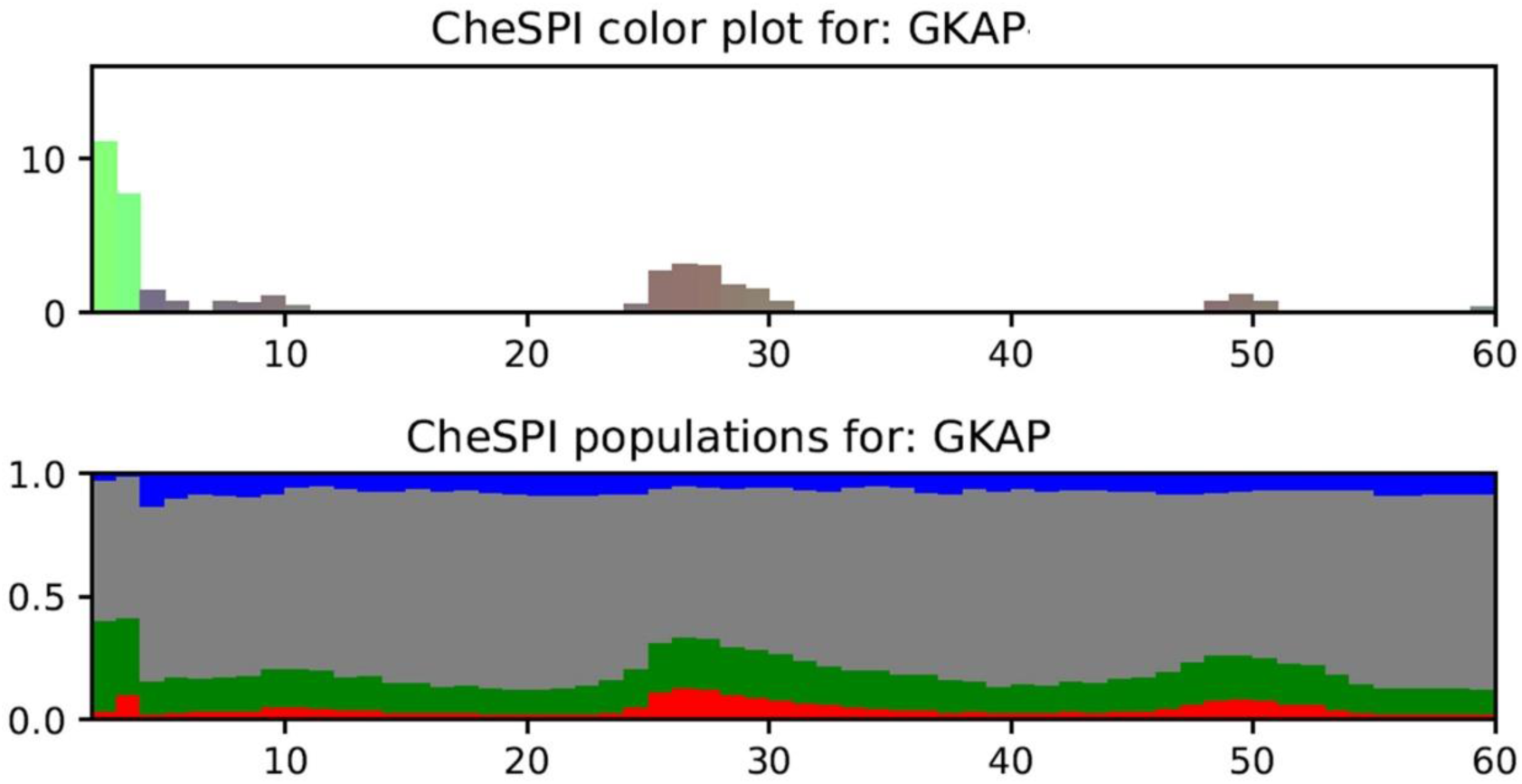
Secondary structure preferences of rGKAP_655-711_ obtained by CheSPI analysis of the assigned chemical shifts [37]. The results are consistent with a largely disordered structure. (A) Bar plot with bar height equal to the CheZOD Z-scores; (B) Accumulated bar-chart of CheSPI secondary structure populations: extended (colored in blue), helical (red), turn (green), and non-folded (grey) local structures.

**Supplementary Figure S4.**
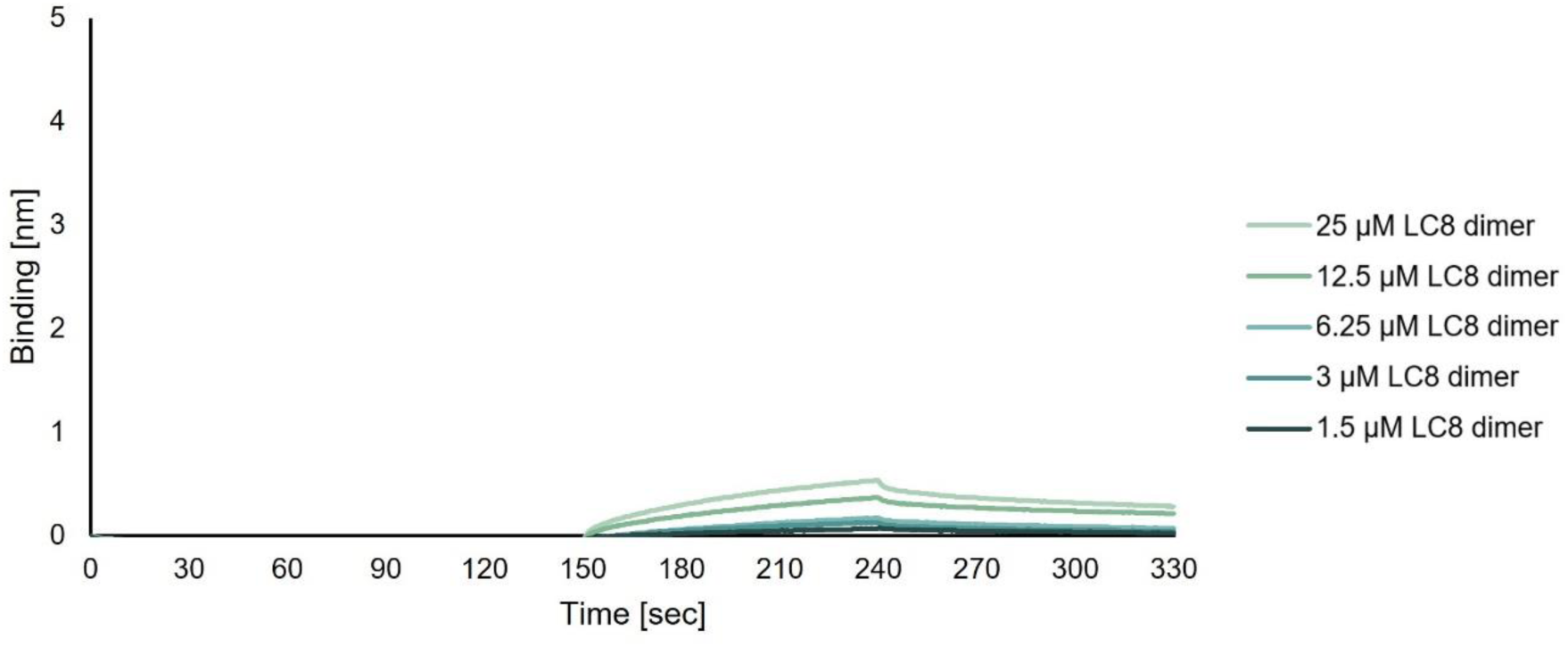
BLI sensogram for the interaction of rGKAP_655-711_ and LC8, control measurement to assess the aspecific binding of LC8 in the absence of GKAP.

**Supplementary Figure S5.**
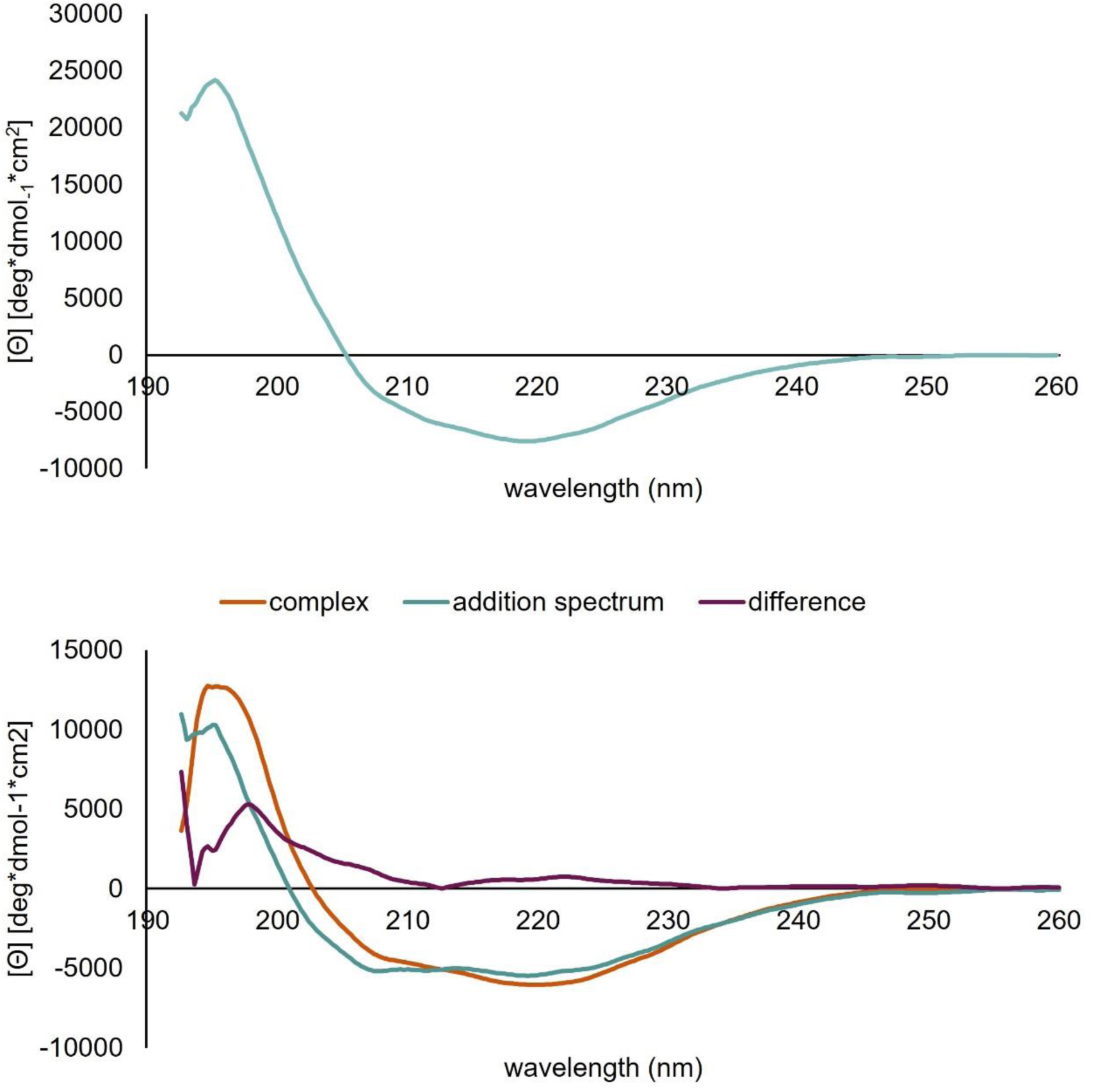
Circular dichroism measurements of the LC8 dimer and the GKAP-LC8 complex. (A) The CD spectrum of LC8 indicates a well-folded structure and matches previously reported spectra [67]. (B) The CD spectrum of the GKAP:LC8 complex is slightly different from the sum of the spectra of the individual partners, consistent with the presence of the interaction and the preservation of the folded LC8 structure. The spectrum is not indicative of any large-scale structural rearrangements.

**Supplementary Figure S6.**
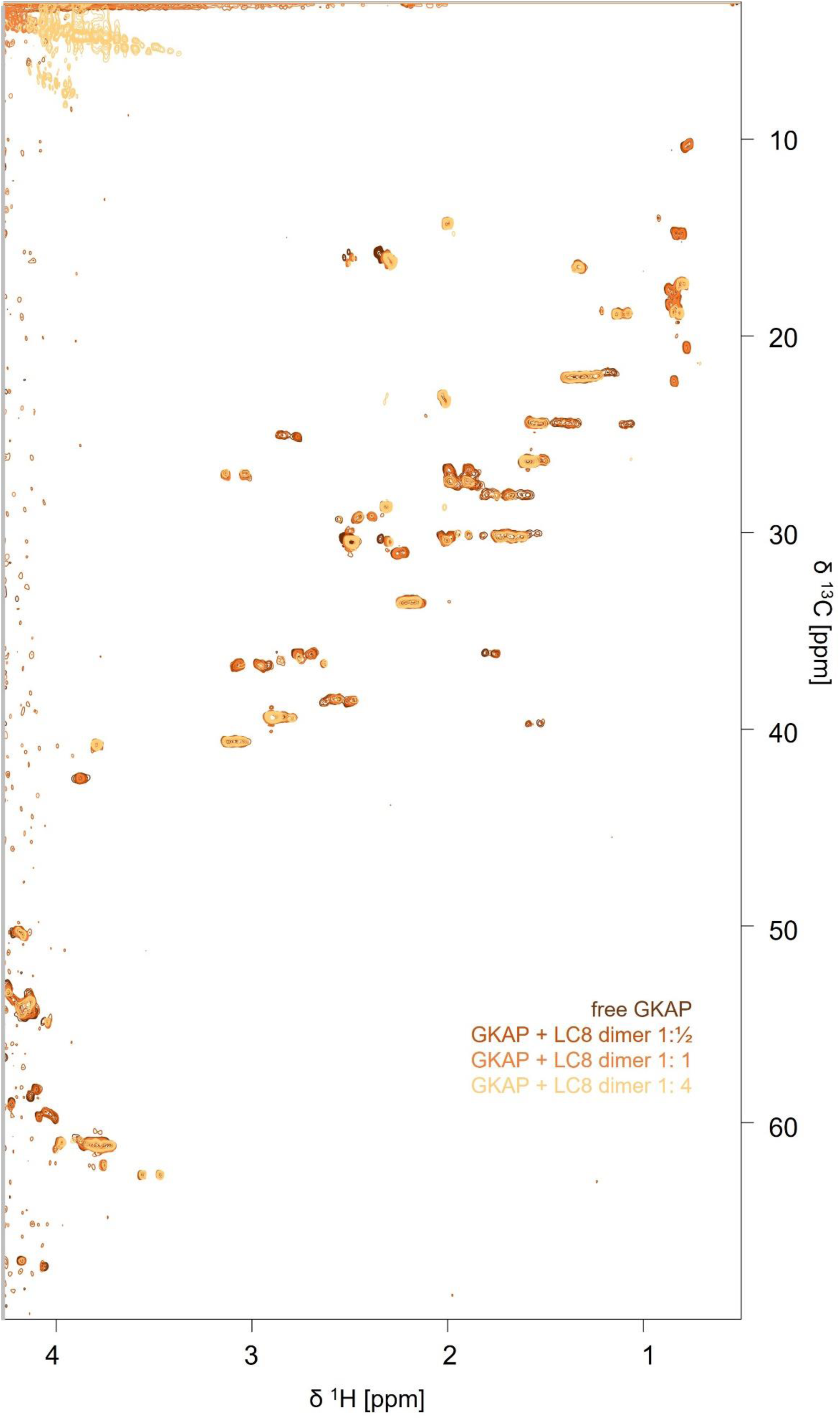
^1^H-^13^C HSQC (aliphatic) of the rGKAP_655-711_ construct titrated with LC8.

**Supplementary Figure S7.**
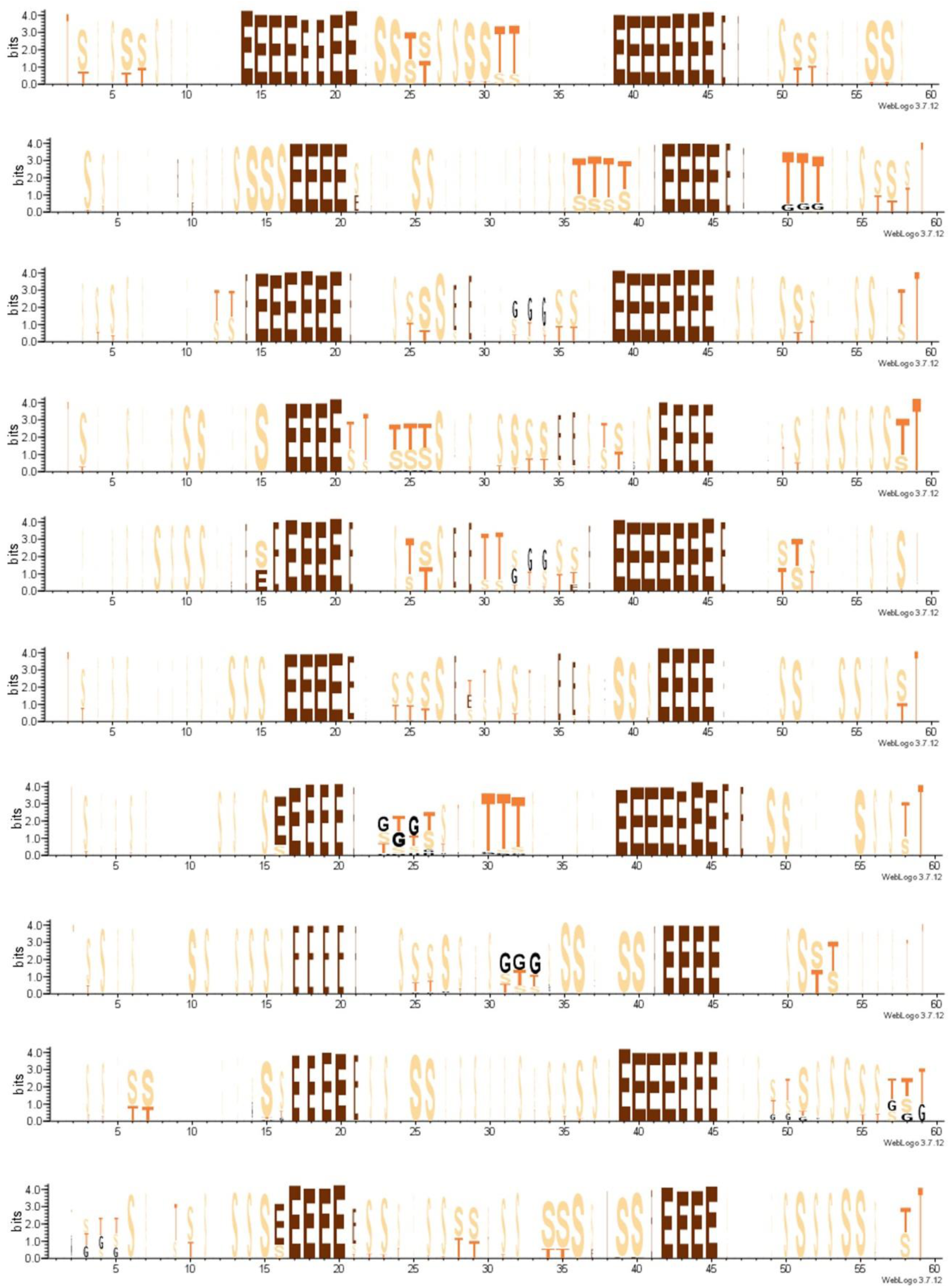
DSSPCont result of the respective GKAP chains (from all 5 simulations, for both GKAP chains)

**Supplementary Figure S8.**
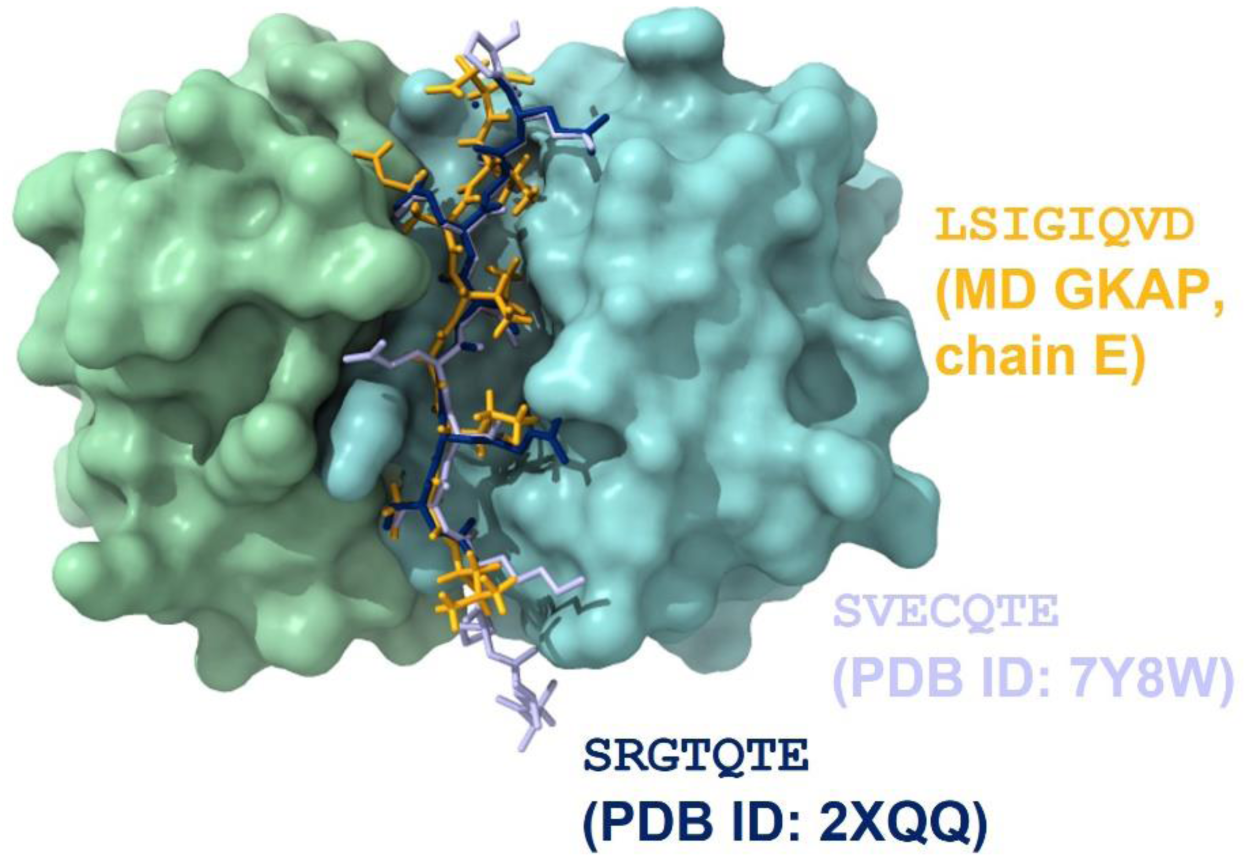
Structural representation of the interaction of the LC8 dimer with peptides containing different binding sequences. Yellow: selected structure from our simulation of the first binding motif of GKAP; Violet: Sao-1 peptide (PDB ID: 7Y8W [51]); Blue: Synthetic peptide based on in vitro evolution results (PDB ID: 2XQQ [14])

**Supplementary Figure S9.**
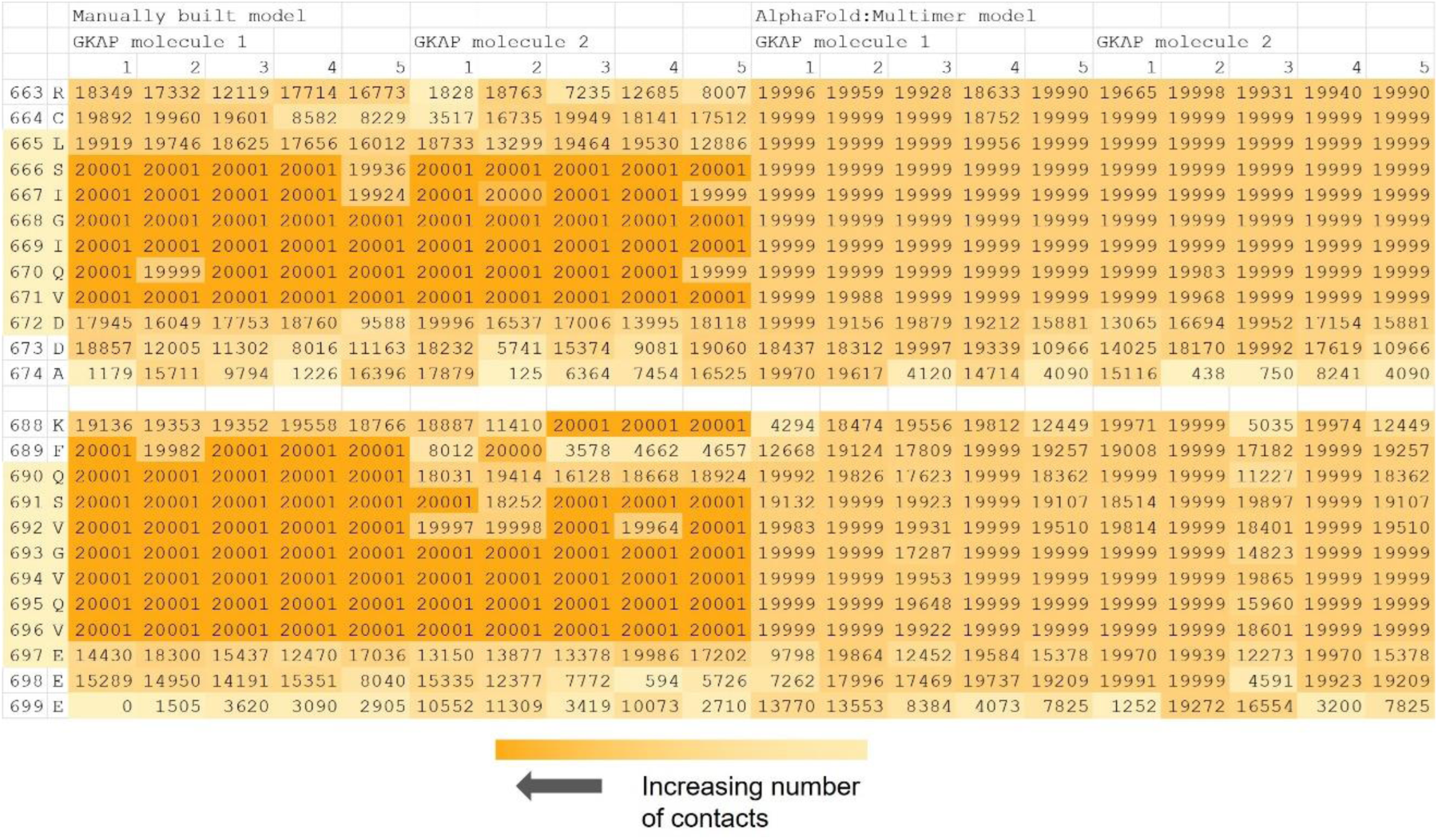
Heatmap of the differences in atomic contacts calculated with Voronota for the two LC8 binding sites of GKAP. Data are shown for both GKAP chains in the manually built models and also for those generated with AlphaFold Multimer. The numbers represent the number of frames in the 200 ns molecular dynamics simulations (20001 frames overall) where the given residue is in contact with another residue on any other polypeptide chain of the complex. For example, F39, located before the second binding motif, is often in contact with one of the LC8 dimers, sometimes with the one at the first binding site.

**Supplementary Figure S10.**
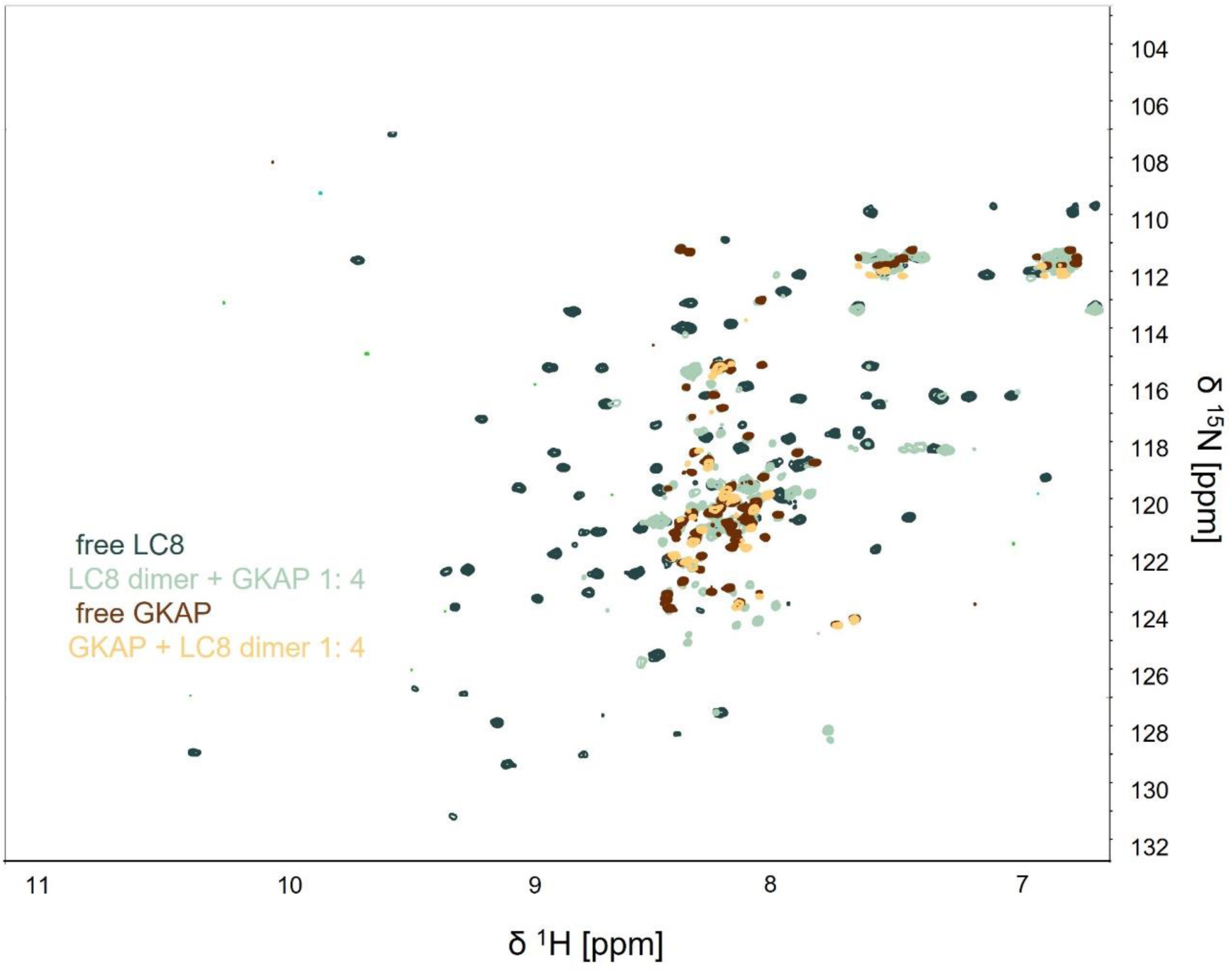
Free LC8 and LC8 titration endpoint overlaid with free GKAP and GKAP titration endpoint. The majority of the free LC8 peaks disappear at the titration endpoint, but many new peaks emerge with ppm values where free GKAP and “forward” GKAP titration endpoint peaks are also visible.

**Supplementary Figure S11.**
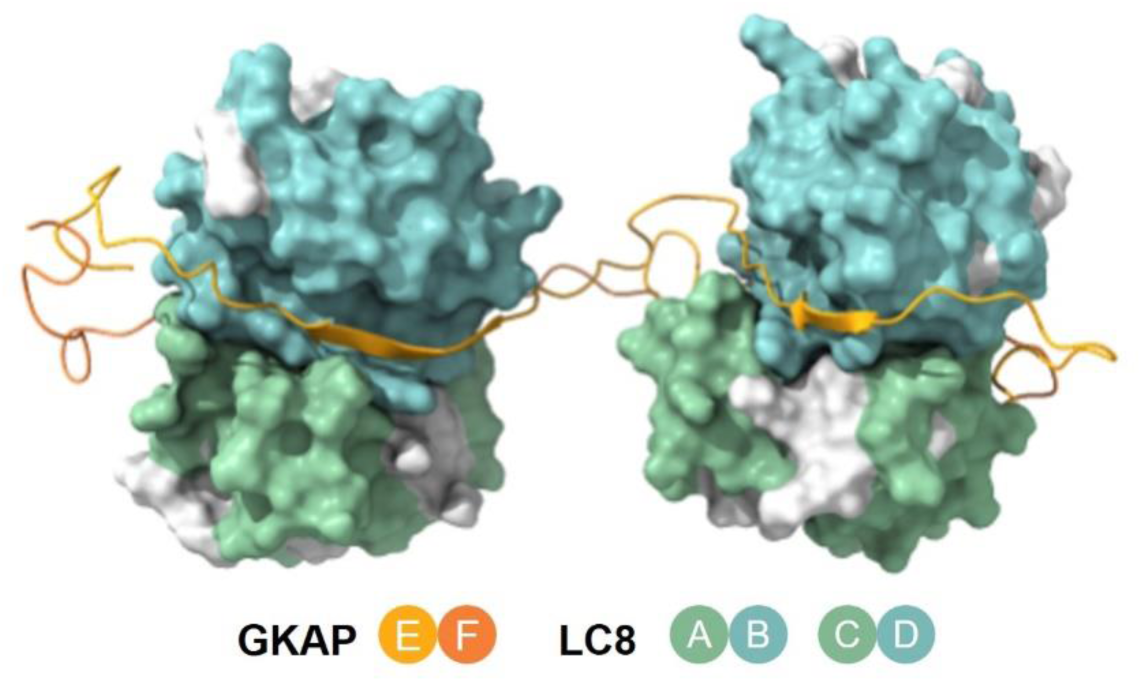
Positions of LC8 residues that are likely not involved in either dimerization or ligand binding. These residues, namely A21, C24, A25, Y41, I42, F46, I74, F76, L85 are shown in gray.

**Supplementary Figure S12.**
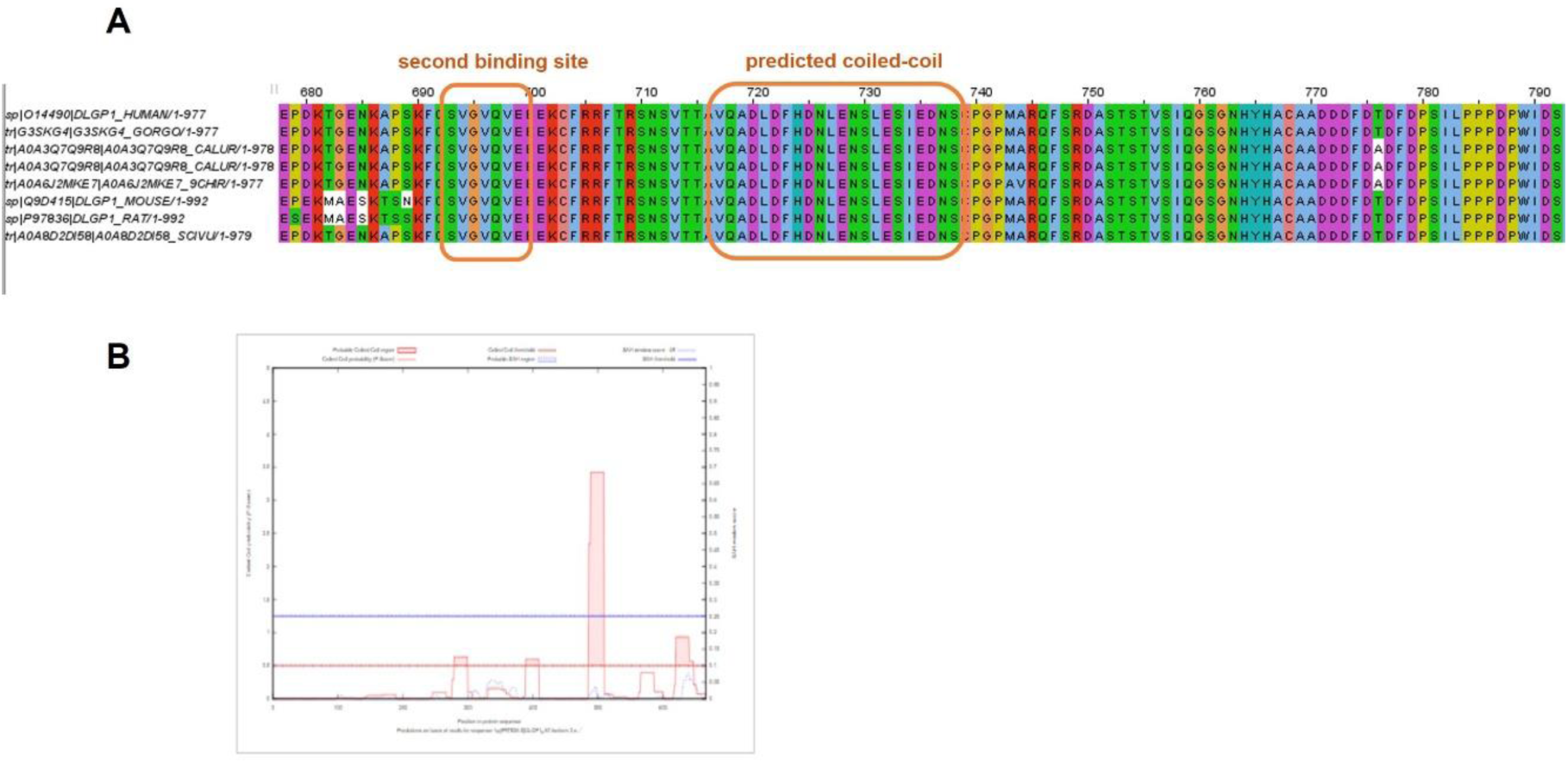
(A) Multiple sequence alignment of selected GKAP sequences with the second binding site and its C-terminal flanking segment, with the binding site and the predicted coiled coil boxed; (B) Prediction output from wagga-wagga with windowsize=21. The prediction could be compatible with the presence of a weak and noncanonical coiled coil in region 722-735.

